## Supplementary material for "All-at-once RNA folding with 3D motif prediction framed by evolutionary information": CsrC

# STOCKHOLM 1.0
#=GF ID CsrC
#=GF AC RF00084
#=GF DE CsrC RNA family
#=GF AU Bateman A; 0000-0002-6982-4660
#=GF GA 61.0
#=GF NC 55.8
#=GF TC 72.6
#=GF PI SraK
#=GF SE Bateman A
#=GF SS Predicted; PFOLD
#=GF TP Gene; sRNA;
#=GF BM cmbuild -F CM SEED
#=GF CB cmcalibrate --mpi CM
#=GF SM cmsearch --cpu 4 --verbose --nohmmonly -T 16.55 -Z 2958934 CM SEQDB
#=GF CL CL00106
#=GF DR SO; 0000370; small_regulatory_ncRNA;
#=GF RN [1]
#=GF RM 12694612
#=GF RT A novel sRNA component of the carbon storage regulatory system of
#=GF RT Escherichia coli.
#=GF RA Weilbacher T, Suzuki K, Dubey AK, Wang X, Gudapaty S, Morozov I, Baker CS,
#=GF RA Georgellis D, Babitzke P, Romeo T
#=GF RL Mol Microbiol 2003;48:657-670.
#=GF RN [2]
#=GF RM 11448770
#=GF RT Novel small RNA-encoding genes in the intergenic regions of Escherichia
#=GF RT coli.
#=GF RA Argaman L, Hershberg R, Vogel J, Bejerano G, Wagner EG, Margalit H,
#=GF RA Altuvia S
#=GF RL Curr Biol 2001;11:941-950.
#=GF CC The 245 nucleotide sRNA of Escherichia coli, CsrC, was discovered using a
#=GF CC genetic screen for factors that regulate glycogen biosynthesis. CsrC RNA
#=GF CC binds multiple copies of CsrA Pfam:PF02599, a protein that
#=GF CC post-transcriptionally regulates central carbon flux, biofilm formation
#=GF CC and motility in E. coli. CsrC antagonises the regulatory effects of CsrA,
#=GF CC presumably by sequestering this protein. The discovery of CsrC is
#=GF CC intriguing, in that a similar sRNA, CsrB, performs essentially the same
#=GF CC function. Both sRNAs possess similar imperfect repeat sequences (18 in
#=GF CC CsrB, nine in CsrC), primarily localised in the loops of predicted
#=GF CC hairpins, which may serve as CsrA binding elements. Transcription of csrC
#=GF CC increases as the culture approaches the stationary phase of growth and is
#=GF CC indirectly activated by CsrA via the response regulator UvrY [1]. This RNA
#=GF CC was also discovered in E. coli during a large scale screen [2]. The gene
#=GF CC called SraK, was highly abundant in stationary phase, but low levels could
#=GF CC be detected in exponentially growing cells as well [2].
#=GF WK CsrC_RNA_family
#=GF SQ 4

CP000647.1/4572620-4572875 CAACGGCGGGGAUGCUGAAAGGAUUAG-GACUC-AGGACGAGGAGUCAGGAAGCGCCAGGAGGCGAAGACCACAGGAUAGUCAGGAUGACGCGCGUCUUGGAGACGUU-UUACAGGGAAAAGGAAGAA-ACAGGGACCGUUGGACGCUAAGGGAAAGACAGGGUGAGUCGUUAGCGAACAGGGAAAGAAGAGCCCGUUAA
CP000653.1/4458195-4457944 CCACGGCGAGGAUGCCAUCAGGAUUA-UGACUUAAGGAUGA-GGGUC-AGGAGCGCCAGGAGGCGAAGACU-CAGGAUUACAAGGACGACA--CGCC-AGGAGGCG-UUUAAAAGGGAAAUGGAACCGUAUCACGGAUGAUUG-CGCUAAAGGAUAAACAGGGUUAGUUGUUGGCGAACAGGGAUUGUAAAACCCGUUUA
CP001127.1/4088494-4088746 AUAG--CGAGGACGCUAACAGGAUCAACGACUC-AGGAUGA-GGGUC-AGGAGCGCCAGGAGGCGAAGACA-CAGGAUUGUCAGGAAGACAAACGUC-CGGAGACGUUAGUAAAAGGAAAUGGAAACA-ACAUGGAAUGUUCCAGGCUAAGGGAAAAACAGGGCGUGUUGAUAGCCAACAGGGAUGGUGGAACCCGUUAA
DQ328981.1/142-396         ACAUAGCGAGGACGCUAACAGGAUCAACGACUC-AGGAUGA-GGGUC-AGGAGCGCCAGGAGGCGAAGACA-CAGGAUUGUCAGGAAGACAAACGUC-CGGAGACGUUAGUAAAAGGAAAUGGAAACA-ACAUGGAAUGUUCCAGGCUAAGGGAAAAACAGGGCGUGUUGAUAGCCAACAGGGAUGGUGGAACCCGUUAA
#=GC SS_cons               ::::<<<<____>>>>:::::::::::<<<<<<._______.>>>>>.>::::<<<<____>>>>::<<<<.______>>>>::::::::::<<<<<.____>>>>>:::::::::::::::::::<<.<<<<<--<<<<<<<________>>>--->>>>-->>>>>>>:::::::<<_______>>:::<<<<_____
#=GC RF                    auAugGCgaGGAcGCcaacAGGAUuAacGaCcc.AGGAuGA.ggGuC.gGgAGCGCCAGGAGGCGAAgacc.CAGGAUggucAGGAaGACaaaCGuC.uGGAGaCGuUauuAaAaGGAAAuGGAAacg.acccgGacuGuuccAcGCUAAgggaaAaaCagGGcgggucguUaGCgAaCAGGGAuaGuagAaCCCGUUaA

CP000647.1/4572620-4572875 GGGUGGUCAGUGA-GGAAAAAGGCGACAGAGUAAUCUGUCGCCUUUUUUCUUUGCUUGCUU
CP000653.1/4458195-4457944 GGGUGGUGAGUCAGGAAAAAAGGCGACAGAUUGCUCUGUCGCCUUUUUUCUUUGCUUGCUU
CP001127.1/4088494-4088746 GGGUCGUGAGUCAAGAAAAAAGGCGGCAGAUUACUCUGUCGCCUUUUUUCUUUGCUUGCUU
DQ328981.1/142-396         GGGUCGUGAGUCAAGAAAAAAGGCGGCAGAUUACUCUGUCGCCUUUUUUCUUUGCUUGCUU
#=GC SS_cons               >>>>::::::::::<<<<<<<<<<<<<<<<____>>>>>>>>>>>>>>>>:::::::::::
#=GC RF                    GGGugGUgAGUcAaGaaaaaaGGCGaCaGauUacuCuGuCGCCuuuuuuCUUUGCUUGCUU
//
