## Supplementary material for "All-at-once RNA folding with 3D motif prediction framed by evolutionary information": SNORD46

# STOCKHOLM 1.0
#=GF ID SNORD46
#=GF AC RF00218
#=GF DE Small nucleolar RNA SNORD46
#=GF AU Eberhardt R; 0000-0001-6152-1369
#=GF GA 40.0
#=GF NC 39.8
#=GF TC 40.0
#=GF PI U40
#=GF SE Griffiths-Jones SR
#=GF SS Predicted; Griffiths-Jones SR
#=GF TP Gene; snRNA; snoRNA; CD-box;
#=GF BM cmbuild -F CM SEED
#=GF CB cmcalibrate --mpi CM
#=GF SM cmsearch --cpu 4 --verbose --nohmmonly -T 22.00 -Z 2958934 CM SEQDB
#=GF CL CL00061
#=GF DR SO; 0000593; C_D_box_snoRNA;
#=GF DR GO; 0006396; RNA processing;
#=GF DR GO; 0005730; nucleolus;
#=GF DR snoopy; Mus_musculus300956;
#=GF DR snoopy; Spermophilus_tridecemlineatus300291;
#=GF RN [1]
#=GF RM 8764399
#=GF RT Intron-encoded, antisense small nucleolar RNAs: the characterization of
#=GF RT nine novel species points to their direct role as guides for the
#=GF RT 2'-O-ribose methylation of rRNAs.
#=GF RA Nicoloso M, Qu LH, Michot B, Bachellerie JP
#=GF RL J Mol Biol 1996;260:178-195.
#=GF RN [2]
#=GF RM 12736310
#=GF RT The high diversity of snoRNAs in plants: identification and comparative
#=GF RT study of 120 snoRNA genes from Oryza sativa.
#=GF RA Chen CL, Liang D, Zhou H, Zhuo M, Chen YQ, Qu LH
#=GF RL Nucleic Acids Res 2003;31:2601-2613.
#=GF CC U40 is a C/D box small nucleolar RNA, also known as snoZ153 in plants. The
#=GF CC human sequence has also been annotated in the sequence databases as U46.
#=GF CC U40 is encoded in intron 2 of the ribosomal protein S8 gene in human, and
#=GF CC is hypothesised to guide methylation of 2'-O-ribose residues on 28S
#=GF CC ribosomal RNA [1]. This family represents vertebrate U40.
#=GF WK Small_nucleolar_RNA_SNORD46
#=GF SQ 5

HQ318014.1/785-888 AACUGGAGUGAUGAAAAAGUAUCCUUAGGCGUGGUUGUGGCCGUCUUGGCCACCUUUGUGCCACUUGCCA-GUGCUAGGACUUGUCAUAGUUACACUGACUGUGU
X96646.1/1-98      --GUAGGGUGAUGAAAAAGAAUCCUUAGGCGUGGUUGUGGCCGUCUUGGUCACCUGUGUGCCACUUGCCA-AUGCAAGGACUUGUCAUAGUUACACUGACU----
Z69628.1/1-100     --GUAGGGUGAUGAAAAAGAAUCCUUAGGCGUGGUUGUGGCCGUCUUGGUCACCUGUGUGCCACUUGCCA-AUGCAAGGACUUGUCAUAGUUACACUGACUGU--
X67247.1/1141-1244 AAGUAGGGUGAUGAAAAAGAAUCCUUAGGCGUGGUUGUGGCCGUCUUGGUCACCUGUGUGCCACUUGCCA-AUGCAAGGACUUGUCAUAGUUACACUGACUGUUG
X94604.1/82-186    AACUAGGGUGAUGAAAACGUAUCCUUAGGCGUGGUUAUGGCCGUCUUGGCCACCUGUACACACACUUGCCAAUGCAAGGACUUGUCAUAGUUACACUGACUGUAA
#=GC SS_cons       <<<<<<________________________________________________________________.____________________________>>>>>>
#=GC RF            aaguaGGGUGAUGAAAAAGaAUCCUUAGGCGUGGUUgUGGCCGUCUUGGuCACCUgUgUgCcAcUUgcCA.AUGCAAGGACUUGUCAUAGUUACACUGACugcuu
//
