## Supplementary material for "All-at-once RNA folding with 3D motif prediction framed by evolutionary information": R-scape_motifs.pl

```
#!/usr/bin/perl -w
# 
#  Rank a R-scape output by the F measure
#
use strict;
use Class::Struct;
use POSIX;

use constant GNUPLOT => '/opt/homebrew/bin/gnuplot';
use lib '/Users/erivas/src/Mysrc/R-scape/scripts';
use FUNCS;

use vars qw ($opt_v $opt_F);  # required if strict used
use Getopt::Std;
getopts ('vF:');

# Print a helpful message if the user provides no input file.
if (!@ARGV) {
    print "usage:  R-scape_motifs.pl [options] <R-scape.out>\n\n";
    print "options:\n";
    print "-v    :  be verbose\n";
    exit;
}
my $file_rscape = shift;
my $file_motifs = shift;

my $seeplots = 0;

my $nmotif_HL = 0;
my $nmotif_BL = 0;
my $nmotif_IL = 0;
my $nmotif_J3 = 0;
my $nmotif_J4 = 0;
my $nmotif_BS = 0;
my $nmotif = 0;
my @motif_name;
my @motif_value;
my @motif_type;
my %motif_idx;
parse_motifs($file_motifs, \$nmotif_HL, \$nmotif_BL, \$nmotif_IL, \$nmotif_J3, \$nmotif_J4, \$nmotif_BS,
	     \$nmotif, \@motif_name, \@motif_value, \@motif_type, \%motif_idx);
printf("number of motifs $nmotif\n");
printf("HL $nmotif_HL\n");
printf("BL $nmotif_BL\n");
printf("IL $nmotif_IL\n");
printf("J3 $nmotif_J3\n");
printf("J4 $nmotif_J4\n");
printf("BS $nmotif_BS\n");
for (my $m = 0; $m < $nmotif; $m ++) {
    printf("%d %s %40s %s\n", $motif_idx{$motif_name[$m]}, $motif_type[$m], $motif_name[$m], $motif_value[$m]);
}
my $dir = "/Users/erivas/writing/papers/R3D/Tables";
my $motifs_tbl = "$dir/Table_R3Dgrm.tex";
motifs_table($motifs_tbl, $nmotif_HL, $nmotif_BL, $nmotif_IL, $nmotif_J3, $nmotif_J4, $nmotif_BS,
	     $nmotif, \@motif_name, \@motif_value, \@motif_type, \%motif_idx);


my $nhotif = 7;
my @hotif_name;
my %hotif_idx;
$hotif_name[0] = "NESTED";
$hotif_name[1] = "PK";
$hotif_name[2] = "TRIPLET";
$hotif_name[3] = "HIGHER";
$hotif_name[4] = "NONWC";
$hotif_name[5] = "SCOV";
$hotif_name[6] = "XCOV";
for (my $h = 0; $h < $nhotif; $h ++) {
    $hotif_idx{$hotif_name[$h]} = $h;
}


my %M_found_cov;
my %M_found;
my $M_found_cov_tot = 0;
my $M_found_tot = 0;
my %nfam_with_M_cov;
my %nfam_with_M;
my %nfam_with_H_cov;
my %nfam_with_H;
my $F_with_M_cov_tot = 0;
my $F_with_M_tot = 0;
my %H_found_cov;
my %H_found;
parse_rscape($file_rscape, $opt_F,
	     $nhotif, \@hotif_name, \%hotif_idx,
	     $nmotif, \@motif_name, \%motif_idx,
	     \%M_found_cov, \%M_found,
	     \$M_found_cov_tot, \$M_found_tot,
	     \%nfam_with_M_cov, \%nfam_with_M,
	     \%nfam_with_H_cov, \%nfam_with_H,
	     \$F_with_M_cov_tot, \$F_with_M_tot,
	     \%H_found_cov, \%H_found,
	     1,
	     $seeplots);

my $motifsfile = ($opt_F)?  "$file_rscape.$opt_F.motifs" : "$file_rscape.motifs";
motifs_write($motifsfile, $opt_F,
	     \%M_found_cov, \%M_found,
	     $M_found_cov_tot, $M_found_tot,
	     \%nfam_with_M_cov, \%nfam_with_M,
	     \%nfam_with_H_cov, \%nfam_with_H,
	     $F_with_M_cov_tot, $F_with_M_tot,
	     \%H_found_cov, \%H_found);


my $SSU = "RF01960";
my %SSU_M_found_cov;
my %SSU_M_found;
my $SSU_M_found_cov_tot = 0;
my $SSU_M_found_tot = 0;
my %SSU_nfam_with_M_cov;
my %SSU_nfam_with_M;
my %SSU_nfam_with_H_cov;
my %SSU_nfam_with_H;
my $SSU_F_with_M_cov_tot = 0;
my $SSU_F_with_M_tot = 0;
my %SSU_H_found_cov;
my %SSU_H_found;
parse_rscape($file_rscape, $SSU,
	     $nhotif, \@hotif_name, \%hotif_idx,
	     $nmotif, \@motif_name, \%motif_idx,
	     \%SSU_M_found_cov, \%SSU_M_found,
	     \$SSU_M_found_cov_tot, \$SSU_M_found_tot,
	     \%SSU_nfam_with_M_cov, \%SSU_nfam_with_M,
	     \%SSU_nfam_with_H_cov, \%SSU_nfam_with_H,
	     \$SSU_F_with_M_cov_tot, \$SSU_F_with_M_tot,
	     \%SSU_H_found_cov, \%SSU_H_found,
	     0,
	     $seeplots);

my $LSU = "RF02543";
my %LSU_M_found_cov;
my %LSU_M_found;
my $LSU_M_found_cov_tot = 0;
my $LSU_M_found_tot = 0;
my %LSU_nfam_with_M_cov;
my %LSU_nfam_with_M;
my %LSU_nfam_with_H_cov;
my %LSU_nfam_with_H;
my $LSU_F_with_M_cov_tot = 0;
my $LSU_F_with_M_tot = 0;
my %LSU_H_found_cov;
my %LSU_H_found;
parse_rscape($file_rscape, $LSU,
	     $nhotif, \@hotif_name, \%hotif_idx,
	     $nmotif, \@motif_name, \%motif_idx,
	     \%LSU_M_found_cov, \%LSU_M_found,
	     \$LSU_M_found_cov_tot, \$LSU_M_found_tot,
	     \%LSU_nfam_with_M_cov, \%LSU_nfam_with_M,
	     \%LSU_nfam_with_H_cov, \%LSU_nfam_with_H,
	     \$LSU_F_with_M_cov_tot, \$LSU_F_with_M_tot,
	     \%LSU_H_found_cov, \%LSU_H_found,
	     0,
	     $seeplots);

$motifsfile = "$file_rscape.together.motifs";
motifs_write_together($motifsfile,
		      \@motif_type, \%motif_idx,
		      \%M_found_cov, \%M_found,
		      $M_found_cov_tot, $M_found_tot,
		      \%nfam_with_M_cov, \%nfam_with_M,
		      \%nfam_with_H_cov, \%nfam_with_H,
		      $F_with_M_cov_tot, $F_with_M_tot,
		      \%H_found_cov, \%H_found,
		      \%SSU_M_found_cov, \%SSU_M_found,
		      $SSU_M_found_cov_tot, $SSU_M_found_tot,
		      \%SSU_H_found_cov, \%SSU_H_found,
		      \%LSU_M_found_cov, \%LSU_M_found,
		      $LSU_M_found_cov_tot, $LSU_M_found_tot,
		      \%LSU_H_found_cov, \%LSU_H_found);


sub parse_rscape{
    my ($rfile, $selectF,
	$nhotif, $ref_hotif_name, $ref_hotif_idx,
	$nmotif, $ref_motif_name, $ref_motif_idx,	
	$ref_M_found_cov, $ref_M_found,	
	$ret_M_found_cov_tot, $ret_M_found_tot,
	$ref_nfam_with_M_cov, $ref_nfam_with_M,
	$ref_nfam_with_H_cov, $ref_nfam_with_H,
	$ret_F_with_M_cov_tot, $ret_F_with_M_tot,
	$ref_H_found_cov, $ref_H_found,
	$do_FamsPerMotif,
	$seeplots) = @_;

    my @HbyF;
    my @HbyF_cov;
    my $nhotif_tot = 0;
    my @HbyF_given;
    my @HbyF_given_cov;
    my $nhotif_given_tot = 0;
    
    my @MbyF;
    my @MbyF_cov;
    my $nmotif_tot = 0;
    my @MbyF_given;
    my @MbyF_given_cov;
    my $nmotif_given_tot = 0;

    my $nHL;
    my $nBL;
    my $nIL;
    my $nJ3;
    my $nJ4;
    my $nBS;
    my $nRM = 0;
    
    my $nHL_tot;
    my $nBL_tot;
    my $nIL_tot;
    my $nJ3_tot;
    my $nJ4_tot;
    my $nBS_tot;
    my $nRM_tot = 0;
   
    my $n_rfam = 0;
    my %rfam_idx;
    my @rfam_name;
    my $rfam;
    my $is_cacofold;
    open (FILE, "$rfile") || die;
    while(<FILE>) {
	# the R3D grammar
	if    (/^HL\s+(\d+)\s*$/) {
	    $nHL     = $1;
	    $nHL_tot = $1;
	    $nRM     += $nHL;
	    $nRM_tot += $nHL_tot;
	}
	elsif (/^BL\s+(\d+)\/(\d+)\s*$/) {
	    $nBL     = $1;
	    $nBL_tot = $2;
	    $nRM     += $nBL;
	    $nRM_tot += $nBL_tot;
	}
	elsif (/^IL\s+(\d+)\/(\d+)\s*$/) {
	    $nIL     = $1;
	    $nIL_tot = $2;
	    $nRM     += $nIL;
	    $nRM_tot += $nIL_tot;
	}
	elsif (/^J3\s+(\d+)\/(\d+)\s*$/) {
	    $nJ3     = $1;
	    $nJ3_tot = $2;
	    $nRM     += $nJ3;
	    $nRM_tot += $nJ3_tot;
	}
	elsif (/^J4\s+(\d+)\/(\d+)\s*$/) {
	    $nJ4     = $1;
	    $nJ4_tot = $2;
	    $nRM     += $nJ4;
	    $nRM_tot += $nJ4_tot; 
	}
	elsif (/^BS\s+(\d+)\s*$/) {
	    $nBS     = $1;
	    $nBS_tot = $1;
	    $nRM     += $nBS;
	    $nRM_tot += $nBS_tot;
	}
	elsif (/^# MSA (\S+)\s+nseq/) {
	    $rfam = $1;
	    
	    if (!$selectF || ($selectF && $rfam =~ /$selectF/)) {
		$rfam_idx{$rfam}    = $n_rfam;
		$rfam_name[$n_rfam] = $rfam;
		#printf("Rfam $rfam idx %d %s\n", $rfam_idx{$rfam}, $rfam_name[$rfam_idx{$rfam}]); 
		$n_rfam ++;
	    }
	}
       }
    close(FILE);
    printf("HL %d/%d\n", $nHL, $nHL_tot);
    printf("BL %d/%d\n", $nBL, $nBL_tot);
    printf("IL %d/%d\n", $nIL, $nIL_tot);
    printf("J3 %d/%d\n", $nJ3, $nJ3_tot);
    printf("J4 %d/%d\n", $nJ4, $nJ4_tot);
    printf("BS %d/%d\n", $nBS, $nBS_tot);
    printf("RM %d/%d\n", $nRM, $nRM_tot);
    if ($nmotif != $nRM) { printf("the number of motifs does not match %d vs %d\n", $nmotif, $nRM); }
    $nmotif_tot = $nRM_tot;
    for (my $r = 0; $r < $nmotif; $r ++) {
	for (my $f = 0; $f < $n_rfam; $f ++) {
	    $MbyF[$r][$f] = 0;
	    $MbyF_cov[$r][$f] = 0;
	    $MbyF_given[$r][$f] = 0;
	    $MbyF_given_cov[$r][$f] = 0;
	}
    }
    
    for (my $h = 0; $h < $nhotif; $h ++) {
	for (my $f = 0; $f < $n_rfam; $f ++) {
	    $HbyF[$h][$f] = 0;
	    $HbyF_cov[$h][$f] = 0;
	    $HbyF_given[$h][$f] = 0;
	    $HbyF_given_cov[$h][$f] = 0;
	}
    }

    my $name;
    my $nbp;
    my $nbp_cov;
    my $hotif_idx;
    
    my $nhl;
    my $nhl_cov;
    my $rfam_idx;
    my $motif_idx;
    my $alen;
    my $nseq;
    my %Rfam_time;
    my %Rfam_alen;
    my %Rfam_nseq;
    my $time;
    my $n_rfam_done = 0;
    my $doit;
    open (FILE, "$rfile") || die;
    while(<FILE>) {
	# the results
	if (/^# MSA (\S+)\s+nseq/) {
	    $rfam = $1;
	    $rfam_idx = $rfam_idx{$rfam};
	    $doit = 1;
	    if ($selectF && !($rfam =~ /$selectF/)) { $doit = 0; }
	}
	elsif ($doit && /^# The predicted CaCoFold structure/) {
	    $is_cacofold = 1;
	}
	elsif ($doit && /# MSA (\S+) alen=(\d+) nseq=(\d+) time=(\S+) secs/) {
	    my $name = $1;
	    $alen = $2;
	    $nseq = $3;
	    $time = $4;
	    $n_rfam_done ++;
	    if ($name =~ /^$rfam$/) {} else { print "$name and $rfam do not match\n"; die; }
	    
	    $Rfam_time{$rfam} = $time;
	    $Rfam_alen{$rfam} = $alen;
	    $Rfam_nseq{$rfam} = $nseq;
	    #printf("%s %d %d %f %f\n", $name, $alen, $nseq, $alen*$nseq, $time);
	    
	    $is_cacofold = 0;
	}
	elsif ($doit && /^# RM_HELIX (\S+)\s+\S+\s+\S+\, nbp = (\d+) nbp_cov = (\d+)\s*$/) {
	    $name    = $1; 
	    $nbp     = $2;
	    $nbp_cov = $3;
	    $hotif_idx = $$ref_hotif_idx{$name};

	    if ($is_cacofold) {
		$HbyF    [$hotif_idx][$rfam_idx] += 1;
		$HbyF_cov[$hotif_idx][$rfam_idx] += ($nbp_cov > 0)? 1:0;

		# join together PK(1)+TRIPLET)2_ into HIGHER_ORDER(3)
		if ($hotif_idx == 1 || $hotif_idx == 2) {
		    $hotif_idx = 3;
		    $HbyF    [$hotif_idx][$rfam_idx] += 1;
		    $HbyF_cov[$hotif_idx][$rfam_idx] += ($nbp_cov > 0)? 1:0;		    
		}

	    }
	    else {
		$HbyF_given    [$hotif_idx][$rfam_idx] += 1;
		$HbyF_given_cov[$hotif_idx][$rfam_idx] += ($nbp_cov > 0)? 1:0;
		
		# join together PK(1)+TRIPLET)2_ into HIGHER_ORDER(3)
		if ($hotif_idx == 1 || $hotif_idx == 2) {
		    $hotif_idx = 3;
		    $HbyF_given    [$hotif_idx][$rfam_idx] += 1;
		    $HbyF_given_cov[$hotif_idx][$rfam_idx] += ($nbp_cov > 0)? 1:0;
		}
	    }
	}
	elsif ($doit && /^# RM_HL (\S+)\s+\S+\, nbp = \d+ nbp_cov = \d+ nhl_b_cov (\d+)\/(\d+)/) {
	    $name    = $1; 
	    $nhl_cov = $2;
	    $nhl     = $3;
	    if ($name =~ /^(\S+)\.rot/) { $name = $1; }
	    $motif_idx = $$ref_motif_idx{$name};
	    
	    if ($is_cacofold) {
		$MbyF    [$motif_idx][$rfam_idx] += 1;
		$MbyF_cov[$motif_idx][$rfam_idx] += ($nhl_cov > 0)? 1:0;
	    }
	    else {
		$MbyF_given    [$motif_idx][$rfam_idx] += 1;
		$MbyF_given_cov[$motif_idx][$rfam_idx] += ($nhl_cov > 0)? 1:0;
	    }
	}
	elsif ($doit && /^# RM_BL (\S+)\s+\S+\s+\S+\, nbp = \d+ nbp_cov = \d+ nhl_b_cov (\d+)\/(\d+)/) {
	    $name    = $1;
	    $nhl_cov = $2;
	    $nhl     = $3;
	    if ($name =~ /^(\S+)\.rot/) { $name = $1; }
	    $motif_idx = $$ref_motif_idx{$name};

	    if ($is_cacofold) {		
		$MbyF    [$motif_idx][$rfam_idx] += 1;
		$MbyF_cov[$motif_idx][$rfam_idx] += ($nhl_cov > 0)? 1:0;
	    }
	    else {
		$MbyF_given    [$motif_idx][$rfam_idx] += 1;
		$MbyF_given_cov[$motif_idx][$rfam_idx] += ($nhl_cov > 0)? 1:0;
	    }
	}
	elsif ($doit && /^# RM_IL (\S+)\s+\S+\s+\S+\, nbp = \d+ nbp_cov = \d+ nhl_b_cov (\d+)\/(\d+)/) {
	    $name    = $1;
	    $nhl_cov = $2;
	    $nhl     = $3;
	    if ($name =~ /^(\S+)\.rot/) { $name = $1; }
	    $motif_idx = $$ref_motif_idx{$name};

	    if ($is_cacofold) {
		$MbyF    [$motif_idx][$rfam_idx] += 1;
		$MbyF_cov[$motif_idx][$rfam_idx] += ($nhl_cov > 0)? 1:0;
	    }
	    else {
		$MbyF_given    [$motif_idx][$rfam_idx] += 1;
		$MbyF_given_cov[$motif_idx][$rfam_idx] += ($nhl_cov > 0)? 1:0;
	    }
	}
	elsif ($doit && /^# RM_J3 (\S+)\s+\S+\s+\S+\s+\S+\, nbp = \d+ nbp_cov = \d+ nhl_b_cov (\d+)\/(\d+)/) {
	    $name    = $1;  
	    $nhl_cov = $2;
	    $nhl     = $3;
	    if ($name =~ /^(\S+)\.rot/) { $name = $1; }	    
	    $motif_idx = $$ref_motif_idx{$name};

	    if ($is_cacofold) {
		$MbyF    [$motif_idx][$rfam_idx] += 1;
		$MbyF_cov[$motif_idx][$rfam_idx] += ($nhl_cov > 0)? 1:0;
	    }
	    else {
		$MbyF_given    [$motif_idx][$rfam_idx] += 1;
		$MbyF_given_cov[$motif_idx][$rfam_idx] += ($nhl_cov > 0)? 1:0;
	    }
	}
	elsif ($doit && /^# RM_J4 (\S+)\s+\S+\s+\S+\s+\S+\s+\S+\, nbp = \d+ nbp_cov = \d+ nhl_b_cov (\d+)\/(\d+)/) {
	    $name    = $1;  
	    $nhl_cov = $2;
	    $nhl     = $3;
	    if ($name =~ /^(\S+)\.rot/) { $name = $1; }	    
	    $motif_idx = $$ref_motif_idx{$name};

	    if ($is_cacofold) {
		$MbyF    [$motif_idx][$rfam_idx] += 1;
		$MbyF_cov[$motif_idx][$rfam_idx] += ($nhl_cov > 0)? 1:0;
	    }
	    else {
		$MbyF_given    [$motif_idx][$rfam_idx] += 1;
		$MbyF_given_cov[$motif_idx][$rfam_idx] += ($nhl_cov > 0)? 1:0;
	    }
	}
	elsif ($doit && $is_cacofold && /^# RM_BS (\S+)\s+\S+\, nbp = \d+ nbp_cov = \d+ nhl_b_cov (\d+)\/(\d+)/) {
	    $name    = $1;
	    $nhl_cov = $2;
	    $nhl     = $3;
	    if ($name =~ /^(\S+)\.rot/) { $name = $1; }
	    $motif_idx = $$ref_motif_idx{$name};

	    if ($is_cacofold) {
		$MbyF    [$motif_idx][$rfam_idx] += 1;
		$MbyF_cov[$motif_idx][$rfam_idx] += ($nhl_cov > 0)? 1:0;
	    }
	    else {
		$MbyF_given    [$motif_idx][$rfam_idx] += 1;
		$MbyF_given_cov[$motif_idx][$rfam_idx] += ($nhl_cov > 0)? 1:0;
	    }
	}
    }
    close (FILE);

    # plot and save the times to file
    #
    my $fam = 0;
    my $timesfile = "$rfile.times";
    open (OUT, ">$timesfile");
    foreach my $name (sort { $Rfam_time{$a} <=> $Rfam_time{$b} } keys %Rfam_time) {
	$fam ++;
	printf(    "%d %f  %s %d %d %f %f\n", $fam, $fam/$n_rfam_done, $name, $Rfam_alen{$name}, $Rfam_nseq{$name}, $Rfam_alen{$name}*$Rfam_nseq{$name}, $Rfam_time{$name});
	printf(OUT "%d %f %s %d %d %f %f\n", $fam, $fam/$n_rfam_done, $name, $Rfam_alen{$name}, $Rfam_nseq{$name}, $Rfam_alen{$name}*$Rfam_nseq{$name}, $Rfam_time{$name});	
    }
    close(OUT);
    plot_times($timesfile, GNUPLOT, $seeplots);
    
    # stats on motifs and families to file
    my %H_found;
    my %H_found_cov;
    my %H_found_given;
    my %H_found_given_cov;
    my %nfam_with_H;
    my %nfam_with_H_cov;
    my %nfam_with_H_given;
    my %nfam_with_H_given_cov;
    my %H_per_fam;
    my %H_per_fam_cov;
    my %H_per_fam_given;
    my %H_per_fam_given_cov;
    for (my $h = 0; $h < $nhotif; $h ++) {
	my $hotif = $$ref_hotif_name[$h];
	$H_found{$hotif}     = 0;
	$H_found_cov{$hotif} = 0;
	$H_found_given{$hotif}     = 0;
	$H_found_given_cov{$hotif} = 0;
	
	$nfam_with_H{$hotif}     = 0;
 	$nfam_with_H_cov{$hotif} = 0;
	$nfam_with_H_given{$hotif}     = 0;
 	$nfam_with_H_given_cov{$hotif} = 0;
    }
    for (my $f = 0; $f < $n_rfam; $f ++) {
	my $fam = $rfam_name[$f];
	$H_per_fam{$fam}     = 0;
	$H_per_fam_cov{$fam} = 0;
 	$H_per_fam_given{$fam}     = 0;
	$H_per_fam_given_cov{$fam} = 0;
    }

     for (my $h = 0; $h < $nhotif; $h ++) {
	my $hotif = $$ref_hotif_name[$h];
	
	for (my $f = 0; $f < $n_rfam_done; $f ++) {
	    my $fam = $rfam_name[$f];

	    $H_found{$hotif}     += $HbyF[$h][$f];
	    $H_found_cov{$hotif} += $HbyF_cov[$h][$f];
	    $H_found_given{$hotif}     += $HbyF_given[$h][$f];
	    $H_found_given_cov{$hotif} += $HbyF_given_cov[$h][$f];
	    
	    $nfam_with_H{$hotif}     += ($HbyF[$h][$f]     > 0)? 1:0;
	    $nfam_with_H_cov{$hotif} += ($HbyF_cov[$h][$f] > 0)? 1:0;
	    $nfam_with_H_given{$hotif}     += ($HbyF_given[$h][$f]     > 0)? 1:0;
	    $nfam_with_H_given_cov{$hotif} += ($HbyF_given_cov[$h][$f] > 0)? 1:0;

	    $H_per_fam{$fam}     += $HbyF[$h][$f];
	    $H_per_fam_cov{$fam} += $HbyF_cov[$h][$f];
	    $H_per_fam_given{$fam}     += $HbyF_given[$h][$f];
	    $H_per_fam_given_cov{$fam} += $HbyF_given_cov[$h][$f];
	}
   }

    my %M_found;
    my %M_found_cov;
    my %M_found_given;
    my %M_found_given_cov;
    my %nfam_with_M;
    my %nfam_with_M_cov;
    my %nfam_with_M_given;
    my %nfam_with_M_given_cov;
    my %M_per_fam;
    my %M_per_fam_cov;
    my %M_per_fam_given;
    my %M_per_fam_given_cov;
    for (my $r = 0; $r < $nmotif; $r ++) {
	my $motif = $$ref_motif_name[$r];
	
	$M_found{$motif}     = 0;
	$M_found_cov{$motif} = 0;
	$M_found_given{$motif}     = 0;
	$M_found_given_cov{$motif} = 0;
	
	$nfam_with_M{$motif}     = 0;
 	$nfam_with_M_cov{$motif} = 0;
 	$nfam_with_M_given{$motif}     = 0;
 	$nfam_with_M_given_cov{$motif} = 0;
    }
    for (my $f = 0; $f < $n_rfam; $f ++) {
	my $fam = $rfam_name[$f];
	$M_per_fam{$fam}     = 0;
	$M_per_fam_cov{$fam} = 0;
	$M_per_fam_given{$fam}     = 0;
	$M_per_fam_given_cov{$fam} = 0;
    }

    my $table_FamsPerMotif_file = "$rfile.FamsPerMotif.tbl";
    if ($do_FamsPerMotif) {
	open (TBL, ">$table_FamsPerMotif_file") || die;
    }
    my $whichmotif = "GAAA_Tetraloop-receptor";
    my $thismotif;
    for (my $r = 0; $r < $nmotif; $r ++) {
	my $motif = $$ref_motif_name[$r];
	$thismotif = 0;
	
	if ($do_FamsPerMotif) { printf(TBL "\nFAMS for MOTIF %s:\n", $motif); }
	
	if ($motif =~ /^$whichmotif$/) {
	    printf("\nFAMS for MOTIF %s:\n", $motif);
	    $thismotif = 1;
	}

	for (my $f = 0; $f < $n_rfam_done; $f ++) {
	    my $fam = $rfam_name[$f];

	    if ($thismotif && $MbyF[$r][$f] > 0) { printf("%s %d(%d)\n", $fam, $MbyF_cov[$r][$f], $MbyF[$r][$f]); }
	    if ($do_FamsPerMotif && $MbyF[$r][$f] > 0) { printf(TBL "%s %d(%d)\n", $fam, $MbyF_cov[$r][$f], $MbyF[$r][$f]); }

	    $M_found{$motif}     += $MbyF[$r][$f];
	    $M_found_cov{$motif} += $MbyF_cov[$r][$f];
	    $M_found_given{$motif}     += $MbyF_given[$r][$f];
	    $M_found_given_cov{$motif} += $MbyF_given_cov[$r][$f];
	    
	    $nfam_with_M{$motif}     += ($MbyF[$r][$f]     > 0)? 1:0;
	    $nfam_with_M_cov{$motif} += ($MbyF_cov[$r][$f] > 0)? 1:0;
	    $nfam_with_M_given{$motif}     += ($MbyF_given[$r][$f]     > 0)? 1:0;
	    $nfam_with_M_given_cov{$motif} += ($MbyF_given_cov[$r][$f] > 0)? 1:0;

	    $M_per_fam{$fam}     += $MbyF[$r][$f];
	    $M_per_fam_cov{$fam} += $MbyF_cov[$r][$f];
	    $M_per_fam_given{$fam}     += $MbyF_given[$r][$f];
	    $M_per_fam_given_cov{$fam} += $MbyF_given_cov[$r][$f];
	}

    }
    if ($do_FamsPerMotif) { close(TBL); }
    
    my $M_found_tot      = 0;
    my $M_found_cov_tot  = 0;
    my $F_with_M_cov_tot = 0;
    my $F_with_M_tot     = 0;
    motifs_stats($selectF,
		\@HbyF,       \@HbyF_cov,
		\@HbyF_given, \@HbyF_given_cov,
		$nhotif, $ref_hotif_name,
		\%H_found,       \%H_found_cov,       \%nfam_with_H,       \%nfam_with_H_cov, 
		\%H_found_given, \%H_found_given_cov, \%nfam_with_H_given, \%nfam_with_H_given_cov,
		$nmotif, $ref_motif_name,
		\%M_found,       \%M_found_cov,       \%nfam_with_M,       \%nfam_with_M_cov, 
		\%M_found_given, \%M_found_given_cov, \%nfam_with_M_given, \%nfam_with_M_given_cov, 
		$n_rfam_done, \@rfam_name, \%rfam_idx,
		\%H_per_fam,     \%H_per_fam_cov,       \%M_per_fam,       \%M_per_fam_cov,
		\%H_per_fam_cov, \%H_per_fam_given_cov, \%M_per_fam_given, \%M_per_fam_given_cov,
		\$M_found_cov_tot, \$M_found_tot, \$F_with_M_cov_tot, \$F_with_M_tot,
		GNUPLOT, $seeplots);

    
    %$ref_M_found_cov      = %M_found_cov;
    %$ref_M_found          = %M_found;	
    $$ret_M_found_cov_tot  = $M_found_cov_tot;
    $$ret_M_found_tot      = $M_found_tot;
    %$ref_nfam_with_M_cov  = %nfam_with_M_cov;
    %$ref_nfam_with_M      = %nfam_with_M;
    %$ref_nfam_with_H_cov  = %nfam_with_H_cov;
    %$ref_nfam_with_H      = %nfam_with_H;
    $$ret_F_with_M_cov_tot = $F_with_M_cov_tot;
    $$ret_F_with_M_tot     = $F_with_M_tot;
    %$ref_H_found_cov      = %H_found_cov;
    %$ref_H_found          = %H_found;

}

sub motifs_stats {
    my ($selectF,
	$ref_HbyF,       $ref_HbyF_cov,
	$ref_HbyF_given, $ref_HbyF_given_cov,
	$nhotif, $ref_hotif_name,
	$ref_H_found,       $ref_H_found_cov,       $ref_nfam_with_H,       $ref_nfam_with_H_cov,
	$ref_H_found_given, $ref_H_found_given_cov, $ref_nfam_with_H_given, $ref_nfam_with_H_given_cov,
	$nmotif, $ref_motif_name,
	$ref_M_found,       $ref_M_found_cov,       $ref_nfam_with_M,       $ref_nfam_with_M_cov,
	$ref_M_found_given, $ref_M_found_given_cov, $ref_nfam_with_M_given, $ref_nfam_with_M_given_cov,
	$n_rfam, $ref_rfam_name, $ref_rfam_idx,
	$ref_H_per_fam,       $ref_H_per_fam_cov,       $ref_M_per_fam,       $ref_M_per_fam_cov,
	$ref_H_per_fam_given, $ref_H_per_fam_given_cov, $ref_M_per_fam_given, $ref_M_per_fam_given_cov,
	$ret_M_found_cov_tot, $ret_M_found_tot, $ret_F_with_M_cov_tot, $ret_F_with_M_tot,
	$gnuplot, $seeplots) = @_;

    my $H_found_tot     = 0;
    my $H_found_cov_tot = 0;   
    foreach my $hotif (reverse sort { $ref_H_found->{$a} <=> $ref_H_found->{$b} } keys %$ref_H_found) {
	$H_found_tot       += $ref_H_found->{$hotif};
	$H_found_cov_tot   += $ref_H_found_cov->{$hotif};

	printf("%30s #Helices %4d/%d #Fam_w_H %4d/%d\n",
	       $hotif, $ref_H_found->{$hotif}, $ref_H_found_cov->{$hotif},
	       $ref_nfam_with_H->{$hotif}, $ref_nfam_with_H_cov->{$hotif});
    }
    printf("Helices found total %d\n", $H_found_tot);
    printf("Helices found with cov support total %d/%d\n\n", $H_found_cov_tot, $H_found_tot);

    my $H_found_given_tot     = 0;
    my $H_found_given_cov_tot = 0;   
    foreach my $hotif (reverse sort { $ref_H_found_given->{$a} <=> $ref_H_found_given->{$b} } keys %$ref_H_found_given) {
	$H_found_given_tot       += $ref_H_found_given->{$hotif};
	$H_found_given_cov_tot   += $ref_H_found_given_cov->{$hotif};
	
	printf("(given) %30s #Helices %4d/%d #Fam_w_H %4d/%d\n",
	       $hotif, $ref_H_found_given->{$hotif}, $ref_H_found_given_cov->{$hotif},
	       $ref_nfam_with_H_given->{$hotif}, $ref_nfam_with_H_given_cov->{$hotif});
    }
    printf("(given) Helices found total %d\n", $H_found_given_tot);
    printf("(given) Helices found with cov support total %d/%d\n\n", $H_found_given_cov_tot, $H_found_given_tot);

    my $M_found_tot     = 0;
    my $M_found_cov_tot = 0;   
    foreach my $motif (reverse sort { $ref_M_found->{$a} <=> $ref_M_found->{$b} } keys %$ref_M_found) {
	$M_found_tot       += $ref_M_found->{$motif};
	$M_found_cov_tot   += $ref_M_found_cov->{$motif};
	
	printf("%30s #Motifs %4d/%d #Fam_w_M %4d/%d\n",
	       $motif, $ref_M_found->{$motif}, $ref_M_found_cov->{$motif},
	       $ref_nfam_with_M->{$motif}, $ref_nfam_with_M_cov->{$motif});
    }
    printf("Motifs found total %d\n", $M_found_tot);
    printf("Motifs found with cov support total %d/%d\n\n", $M_found_cov_tot, $M_found_tot);

    my $M_found_given_tot     = 0;
    my $M_found_given_cov_tot = 0;   
    foreach my $motif (reverse sort { $ref_M_found_given->{$a} <=> $ref_M_found_given->{$b} } keys %$ref_M_found_given) {
	$M_found_given_tot       += $ref_M_found_given->{$motif};
	$M_found_given_cov_tot   += $ref_M_found_given_cov->{$motif};
	
	#printf("(given) %30s #Motifs %4d/%d #Fam_w_M %4d/%d\n",
	#       $motif, $ref_M_found_given->{$motif}, $ref_M_found_given_cov->{$motif},
	#       $ref_nfam_with_M_given->{$motif}, $ref_nfam_with_M_given_cov->{$motif});
    }
    printf("(given) Motifs found total %d\n", $M_found_given_tot);
    printf("(given) Motifs found with cov support total %d/%d\n\n", $M_found_given_cov_tot, $M_found_given_tot);

    my $F_with_H_tot     = 0;
    my $F_with_H_cov_tot = 0;
    foreach my $fam (reverse sort { $ref_H_per_fam->{$a} <=> $ref_H_per_fam->{$b} } keys %$ref_H_per_fam) {
	my $fam_idx = $ref_rfam_idx->{$fam};
	
	$F_with_H_tot     += ($ref_H_per_fam->{$fam} > 0)?     1:0;
	$F_with_H_cov_tot += ($ref_H_per_fam_cov->{$fam} > 0)? 1:0;
	
	#printf("%30s #H %d/%d #NESTED %d/%d #PK %d/%d #TRIPLET %d/%d #HIGHER ORDER %d/%d #MOTIFS %d/%d\n", $fam,
	#       $ref_H_per_fam->{$fam}, $ref_H_per_fam_cov->{$fam},
	#       $ref_HbyF->[0][$fam_idx], $ref_HbyF_cov->[0][$fam_idx], 
	#       $ref_HbyF->[1][$fam_idx], $ref_HbyF_cov->[1][$fam_idx], 
	#       $ref_HbyF->[2][$fam_idx], $ref_HbyF_cov->[2][$fam_idx],
	#       $ref_HbyF->[3][$fam_idx], $ref_HbyF_cov->[3][$fam_idx],
	#       $ref_M_per_fam->{$fam}, $ref_M_per_fam_cov->{$fam});
    }
    my $F_with_H_given_tot     = 0;
    my $F_with_H_given_cov_tot = 0;
    foreach my $fam (reverse sort { $ref_H_per_fam_given->{$a} <=> $ref_H_per_fam_given->{$b} } keys %$ref_H_per_fam_given) {
	my $fam_idx = $ref_rfam_idx->{$fam};
	
	$F_with_H_given_tot     += ($ref_H_per_fam_given->{$fam} > 0)?     1:0;
	$F_with_H_given_cov_tot += ($ref_H_per_fam_given_cov->{$fam} > 0)? 1:0;
	
	#printf("(given) %30s #H %d/%d #NESTED %d/%d #PK %d/%d #TRIPLET %d/%d #HIGHER ORDER %d/%d #MOTIFS %d/%d\n", $fam,
	#       $ref_H_per_fam_given->{$fam}, $ref_H_per_fam_given_cov->{$fam},
	#       $ref_HbyF_given->[0][$fam_idx], $ref_HbyF_given_cov->[0][$fam_idx], 
	#       $ref_HbyF_given->[1][$fam_idx], $ref_HbyF_given_cov->[1][$fam_idx], 
	#       $ref_HbyF_given->[2][$fam_idx], $ref_HbyF_given_cov->[2][$fam_idx],
	#       $ref_HbyF_given->[3][$fam_idx], $ref_HbyF_given_cov->[3][$fam_idx],
	#       $ref_M_per_fam_given->{$fam}, $ref_M_per_fam_given_cov->{$fam});
    }
    printf("Rfam families %d\n", $n_rfam);
    printf("Rfam families with H %d/%d (given) %d/%d\n",
	   $F_with_H_tot, $n_rfam, $F_with_H_given_tot, $n_rfam);
    printf("Rfam families with H cov %d/%d (given) %d/%d\n",
	   $F_with_H_cov_tot, $F_with_H_tot, $F_with_H_given_cov_tot, $F_with_H_given_tot);
    
    
    my $F_with_M_tot     = 0;
    my $F_with_M_cov_tot = 0;
    foreach my $fam (reverse sort { $ref_M_per_fam->{$a} <=> $ref_M_per_fam->{$b} } keys %$ref_M_per_fam) {
	$F_with_M_tot     += ($ref_M_per_fam->{$fam} > 0)?     1:0;
	$F_with_M_cov_tot += ($ref_M_per_fam_cov->{$fam} > 0)? 1:0;
    }
    my $F_with_M_given_tot     = 0;
    my $F_with_M_given_cov_tot = 0;
    foreach my $fam (reverse sort { $ref_M_per_fam_given->{$a} <=> $ref_M_per_fam_given->{$b} } keys %$ref_M_per_fam_given) {
	$F_with_M_given_tot     += ($ref_M_per_fam_given->{$fam} > 0)?     1:0;
	$F_with_M_given_cov_tot += ($ref_M_per_fam_given_cov->{$fam} > 0)? 1:0;
    }
    printf("Rfam families with M %d/%d (given) %d/%d\n",
	   $F_with_M_tot, $n_rfam, $F_with_M_given_tot, $n_rfam);
    printf("Rfam families with M cov %d/%d (given) %d/%d\n",
	   $F_with_M_cov_tot, $F_with_M_tot, $F_with_M_given_cov_tot, $F_with_M_given_tot);


    # together
    foreach my $fam (reverse sort { $ref_H_per_fam->{$a} <=> $ref_H_per_fam->{$b} } keys %$ref_H_per_fam) {
	my $fam_idx = $ref_rfam_idx->{$fam};
	printf("%40s #H %d/%d \(%d/%d\) #NESTED %d/%d \(%d/%d\) #PK %d/%d \(%d/%d\) #TRIPLET %d/%d \(%d/%d\) #HIGHER ORDER %d/%d \(%d/%d\) #MOTIFS %d/%d \(%d/%d\)\n",
	       $fam,
	       $ref_H_per_fam->{$fam},         $ref_H_per_fam_cov->{$fam},
	       $ref_H_per_fam_given->{$fam},   $ref_H_per_fam_given_cov->{$fam},
	       $ref_HbyF->[0][$fam_idx],       $ref_HbyF_cov->[0][$fam_idx], 
	       $ref_HbyF_given->[0][$fam_idx], $ref_HbyF_given_cov->[0][$fam_idx],
	       $ref_HbyF->[1][$fam_idx],       $ref_HbyF_cov->[1][$fam_idx], 
	       $ref_HbyF_given->[1][$fam_idx], $ref_HbyF_given_cov->[1][$fam_idx],
	       $ref_HbyF->[2][$fam_idx],       $ref_HbyF_cov->[2][$fam_idx],
	       $ref_HbyF_given->[2][$fam_idx], $ref_HbyF_given_cov->[2][$fam_idx],
	       $ref_HbyF->[3][$fam_idx],       $ref_HbyF_cov->[3][$fam_idx],
	       $ref_HbyF_given->[3][$fam_idx], $ref_HbyF_given_cov->[3][$fam_idx],
	       $ref_M_per_fam->{$fam},         $ref_M_per_fam_cov->{$fam},
	       $ref_M_per_fam_given->{$fam},   $ref_M_per_fam_given_cov->{$fam}); 	
    }

    foreach my $fam (reverse sort { $ref_M_per_fam->{$a} <=> $ref_M_per_fam->{$b} } keys %$ref_M_per_fam) {
	printf("%30s #M %d/%d\n", $fam, $ref_M_per_fam->{$fam}, $ref_M_per_fam_cov->{$fam});
    }
    #foreach my $fam (reverse sort { $ref_M_per_fam_given->{$a} <=> $ref_M_per_fam_given->{$b} } keys %$ref_M_per_fam_given) {
    #	printf("(given) %30s #M %d/%d\n", $fam, $ref_M_per_fam_given->{$fam}, $ref_M_per_fam_given_cov->{$fam});
    #}


    $$ret_M_found_tot      = $M_found_tot;
    $$ret_M_found_cov_tot  = $M_found_cov_tot;
    $$ret_F_with_M_cov_tot = $F_with_M_cov_tot;
    $$ret_F_with_M_tot     = $F_with_M_tot;
 }

sub motifs_write {
    my ($file, $selectF,
	$ref_M_found_cov, $ref_M_found,
	$M_found_cov_tot, $M_found_tot,
	$ref_nfam_with_M_cov, $ref_nfam_with_M,
	$ref_nfam_with_H_cov, $ref_nfam_with_H,
	$F_with_M_cov_tot, $F_with_M_tot,
	$ref_H_found_cov, $ref_H_found) = @_;
    
    open (OUT, ">$file");
    foreach my $motif (reverse sort { $ref_M_found_cov->{$a} <=> $ref_M_found_cov->{$b} } keys %$ref_M_found_cov) {
	if ($ref_M_found->{$motif} > 0) {
	    if ($selectF) {
		printf("%30s & %d \(%d\) \n",
		       $motif,
		       $ref_M_found_cov->{$motif},     $ref_M_found->{$motif});
		printf(OUT "%30s & %d \(%d\)\n",
		       $motif,
		       $ref_M_found_cov->{$motif},     $ref_M_found->{$motif});
	    }
	    else {
		printf("%30s & %d \(%d\) &  %d \(%d\)\n",
		       $motif,
		       $ref_M_found_cov->{$motif},     $ref_M_found->{$motif}, 
		       $ref_nfam_with_M_cov->{$motif}, $ref_nfam_with_M->{$motif});
		printf(OUT "%30s & %d \(%d\) &  %d \(%d\)\n",
		       $motif,
		       $ref_M_found_cov->{$motif},     $ref_M_found->{$motif}, 
		       $ref_nfam_with_M_cov->{$motif}, $ref_nfam_with_M->{$motif});
	    }
	}
    }
    if ($selectF) {
	printf("Motifs totals & %d \(%d\) \n",
	       $M_found_cov_tot,  $M_found_tot);
	printf(OUT "Motifs totals & %d \(%d\) \n",
	       $M_found_cov_tot,  $M_found_tot);
    }
    else {
	printf("Motifs totals & %d \(%d\) &  %d \(%d\)\n",
	       $M_found_cov_tot,  $M_found_tot, 
	       $F_with_M_cov_tot, $F_with_M_tot );
	printf(OUT "Motifs totals & %d \(%d\) &  %d \(%d\)\n",
	       $M_found_cov_tot,  $M_found_tot, 
	       $F_with_M_cov_tot, $F_with_M_tot );
    }
    
    
    foreach my $hotif (reverse sort { $ref_H_found_cov->{$a} <=> $ref_H_found_cov->{$b} } keys %$ref_H_found_cov) {
	if ($hotif =~ /NESTED/) {
	    if ($selectF) {
		printf("Nested helices & %d \(%d\)\n",
		       $ref_H_found_cov->{$hotif},     $ref_H_found->{$hotif});
		printf(OUT "Nested helices & %d \(%d\)\n",
		       $ref_H_found_cov->{$hotif},     $ref_H_found->{$hotif});
	    }
	    else {
		printf("Nested helices & %d \(%d\) &  %d \(%d\)\n",
		       $ref_H_found_cov->{$hotif},     $ref_H_found->{$hotif}, 
		       $ref_nfam_with_H_cov->{$hotif}, $ref_nfam_with_H->{$hotif});
		printf(OUT "Nested helices & %d \(%d\) &  %d \(%d\)\n",
		       $ref_H_found_cov->{$hotif},     $ref_H_found->{$hotif}, 
		       $ref_nfam_with_H_cov->{$hotif}, $ref_nfam_with_H->{$hotif});
	    }
	}
	if ($hotif =~ /HIGHER/) {
	    if ($selectF) {
		printf(    "PK + higher order & %d \(%d\)\n",
			   $ref_H_found_cov->{$hotif},     $ref_H_found->{$hotif});
		printf(OUT "PK + higher order & %d \(%d\)\n",
		       $ref_H_found_cov->{$hotif},     $ref_H_found->{$hotif});
	    }
	    else {
		printf(    "PK + higher order & %d \(%d\) &  %d \(%d\)\n",
			   $ref_H_found_cov->{$hotif},     $ref_H_found->{$hotif}, 
			   $ref_nfam_with_H_cov->{$hotif}, $ref_nfam_with_H->{$hotif});
		printf(OUT "PK + higher order & %d \(%d\) &  %d \(%d\)\n",
		       $ref_H_found_cov->{$hotif},     $ref_H_found->{$hotif}, 
		       $ref_nfam_with_H_cov->{$hotif}, $ref_nfam_with_H->{$hotif});
	    }
	}
    }
    close(OUT);
    
}


sub motifs_write_together {
    my ($file,
	$ref_motif_type, $ref_motif_idx,
	$ref_M_found_cov, $ref_M_found,
	$M_found_cov_tot, $M_found_tot,
	$ref_nfam_with_M_cov, $ref_nfam_with_M,
	$ref_nfam_with_H_cov, $ref_nfam_with_H,
	$F_with_M_cov_tot, $F_with_M_tot,
	$ref_H_found_cov, $ref_H_found,	
	$ref_SSU_M_found_cov, $ref_SSU_M_found,
	$SSU_M_found_cov_tot, $SSU_M_found_tot,
	$ref_SSU_H_found_cov, $ref_SSU_H_found,
	$ref_LSU_M_found_cov, $ref_LSU_M_found,
	$LSU_M_found_cov_tot, $LSU_M_found_tot,
	$ref_LSU_H_found_cov, $ref_LSU_H_found) = @_;
    
    open (OUT, ">$file");
    foreach my $motif (reverse sort { $ref_M_found_cov->{$a} <=> $ref_M_found_cov->{$b} } keys %$ref_M_found_cov) {
	my $name = $motif;
	$name =~ s/\_/\\\_/g;
	$name =~ s/\-/\\\_/g;
	$name =~ s/\^/\\\_/g;
	my $type = $ref_motif_type->[$ref_motif_idx->{$motif}];
	
	if ($ref_M_found->{$motif} > 0) {
	    printf(    "%s & %30s & %d \(%d\) &  %d \(%d\) &  %d \(%d\) &  %d \(%d\)\\\\ \n",
		   $type, $name,
		   $ref_M_found_cov->{$motif},     $ref_M_found->{$motif}, 
		   $ref_nfam_with_M_cov->{$motif}, $ref_nfam_with_M->{$motif},
		   $ref_SSU_M_found_cov->{$motif}, $ref_SSU_M_found->{$motif},
		   $ref_LSU_M_found_cov->{$motif}, $ref_LSU_M_found->{$motif});
	    printf(OUT "%s & %30s & %d \(%d\) &  %d \(%d\) &  %d \(%d\) &  %d \(%d\)\\\\ \n",
		   $type, $name,
		   $ref_M_found_cov->{$motif},     $ref_M_found->{$motif}, 
		   $ref_nfam_with_M_cov->{$motif}, $ref_nfam_with_M->{$motif},
		   $ref_SSU_M_found_cov->{$motif}, $ref_SSU_M_found->{$motif},
		   $ref_LSU_M_found_cov->{$motif}, $ref_LSU_M_found->{$motif});
	}
    }

   
    printf(    "\\multicolumn\{2\}\{|l|\}\{\\textbf\{3D Motifs all}\} & %d \(%d\) &  %d \(%d\) &  %d \(%d\) &  %d \(%d\)\\\\\n",
	   $M_found_cov_tot,     $M_found_tot, 
	   $F_with_M_cov_tot,    $F_with_M_tot,
	   $SSU_M_found_cov_tot, $SSU_M_found_tot,
	   $LSU_M_found_cov_tot, $LSU_M_found_tot);
   printf(OUT "\\multicolumn\{2\}\{|l|\}\{\\textbf\{3D Motifs all\}\} & %d \(%d\) &  %d \(%d\) &  %d \(%d\) &  %d \(%d\)\\\\\n",
	   $M_found_cov_tot,     $M_found_tot, 
	   $F_with_M_cov_tot,    $F_with_M_tot,
	   $SSU_M_found_cov_tot, $SSU_M_found_tot,
	   $LSU_M_found_cov_tot, $LSU_M_found_tot);
    
    foreach my $hotif (reverse sort { $ref_H_found_cov->{$a} <=> $ref_H_found_cov->{$b} } keys %$ref_H_found_cov) {
	if ($hotif =~ /NESTED/) {
	    printf(    "\\multicolumn\{2\}\{|l|\}\{\\textbf\{Nested helices\}\} & %d \(%d\) &  %d \(%d\) &  %d \(%d\) &  %d \(%d\)\\\\\n",
		   $ref_H_found_cov->{$hotif},     $ref_H_found->{$hotif}, 
		   $ref_nfam_with_H_cov->{$hotif}, $ref_nfam_with_H->{$hotif},
		   $ref_SSU_H_found_cov->{$hotif}, $ref_SSU_H_found->{$hotif},
		   $ref_LSU_H_found_cov->{$hotif}, $ref_LSU_H_found->{$hotif});
	    printf(OUT "\\multicolumn\{2\}\{|l|\}\{\\textbf\{Nested helices\}\} & %d \(%d\) &  %d \(%d\) &  %d \(%d\) &  %d \(%d\)\\\\\n",
		   $ref_H_found_cov->{$hotif},     $ref_H_found->{$hotif}, 
		   $ref_nfam_with_H_cov->{$hotif}, $ref_nfam_with_H->{$hotif},
		   $ref_SSU_H_found_cov->{$hotif}, $ref_SSU_H_found->{$hotif},
		   $ref_LSU_H_found_cov->{$hotif}, $ref_LSU_H_found->{$hotif});
	}
	if ($hotif =~ /HIGHER/) {
	    printf(    "\\multicolumn\{2\}\{|l|\}\{\\textbf\{PK + higher order\}\} & %d \(%d\) &  %d \(%d\)  &  %d \(%d\) &  %d \(%d\)\\\\\n",
		   $ref_H_found_cov->{$hotif},     $ref_H_found->{$hotif}, 
		   $ref_nfam_with_H_cov->{$hotif}, $ref_nfam_with_H->{$hotif},
		   $ref_SSU_H_found_cov->{$hotif}, $ref_SSU_H_found->{$hotif},
		   $ref_LSU_H_found_cov->{$hotif}, $ref_LSU_H_found->{$hotif});
	    printf(OUT "\\multicolumn\{2\}\{|l|\}\{\\textbf\{PK + higher order\}\} & %d \(%d\) &  %d \(%d\)  &  %d \(%d\) &  %d \(%d\)\\\\\n",
		   $ref_H_found_cov->{$hotif},     $ref_H_found->{$hotif}, 
		   $ref_nfam_with_H_cov->{$hotif}, $ref_nfam_with_H->{$hotif},
		   $ref_SSU_H_found_cov->{$hotif}, $ref_SSU_H_found->{$hotif},
		   $ref_LSU_H_found_cov->{$hotif}, $ref_LSU_H_found->{$hotif});
	}
    }

    close(OUT);
    
}

sub plot_times {
    my ($file, $gnuplot, $seeplots) = @_;
    
    my $psfile = "$file.ps";
    #if ($psfile =~ /\/([^\/]+)\s*$/) { $psfile = "$1"; }
    my $pdffile = $psfile;
    if ($pdffile =~ /^(\S+).ps$/) { $pdffile = "$1.pdf"; }

    my $xlabel = "len * nseq";
    my $title  = "";
    
    open(GP,'|'."$gnuplot") || die "Gnuplot: $!";
    
    print GP "set terminal postscript color solid 14\n";
    FUNCS::gnuplot_define_styles (*GP);

    print GP "set output '$psfile'\n";
    
    #print GP "set title \"$title\\n\\n$key\"\n";
    print GP "set title '$title'\n";

    my $cmd = "'$file' using 4:7  title 'alen' ls 1"; 
    print GP "plot $cmd\n";
    $cmd = "'$file' using 7:2  title 'cummulative' ls 1"; 
    print GP "plot $cmd\n";
    $cmd = "'$file' using 5:7  title 'nseq' ls 1"; 
    print GP "plot $cmd\n";
    $cmd = "'$file' using 6:7  title 'alen*nseq' ls 1"; 
    print GP "plot $cmd\n";

    close (GP);

    system ("ps2pdf $psfile $pdffile\n"); 
    system("/bin/rm  $psfile\n");
    if ($seeplots) { system ("open $pdffile&\n"); }
 }


sub parse_motifs {
    my ($file,
	$ret_nmotif_HL, $ret_nmotif_BL, $ret_nmotif_IL, $ret_nmotif_J3, $ret_nmotif_J4, $ret_nmotif_BS,
	$ret_nmotif, $ref_motif_name, $ref_motif_value, $ref_motif_type, $ref_motif_idx) = @_;

    my $nmotif    = 0;
    my $nmotif_HL = 0;
    my $nmotif_BL = 0;
    my $nmotif_IL = 0;
    my $nmotif_J3 = 0;
    my $nmotif_J4 = 0;
    my $nmotif_BS = 0;

    my $name;
    my $value;
    my $type;
    open (FILE, "$file") || die;
    while(<FILE>) {
	if (/^\s*#/) {
	}
	elsif (/^\s*(HL)\s+(\S+\s+\S+\s+\S+)\s+(\S+)\s*$/ || /^\s*(HL)\s+(\S+\s+\S+\s+\S+)\s+(\S+)\s+#/) {
	    $type  = $1;
	    $value = $2;
	    $name  = $3;

	    my $newvalue = $value;
	    
	    $ref_motif_name->[$nmotif]  = $name;
	    $ref_motif_value->[$nmotif] = $newvalue;
	    $ref_motif_type->[$nmotif]  = $type;
	    $ref_motif_idx->{$name}     = $nmotif;
	    $nmotif    += 1;
	    $nmotif_HL += 1;
	}
	elsif (/^\s*(BL)\s+(\S+\s+\S+\s+\S+)\s+(\S+)\s*$/ || /^\s*(BL)\s+(\S+\s+\S+\s+\S+)\s+(\S+)\s+#/) {
	    $type  = $1;
	    $value = $2;
	    $name  = $3;

	    my $newvalue = $value;
	    
	    $ref_motif_name->[$nmotif]  = $name;
	    $ref_motif_value->[$nmotif] = $newvalue;
	    $ref_motif_type->[$nmotif]  = $type;
	    $ref_motif_idx->{$name}     = $nmotif;
	    $nmotif    += 1;
	    $nmotif_BL += 1;
	}
	elsif (/^\s*(IL)\s+(\S+\s+\S+\s+\S+\s+\S+\s+\S+\s+\S+)\s+(\S+)\s*$/ || /^\s*(IL)\s+(\S+\s+\S+\s+\S+\s+\S+\s+\S+\s+\S+)\s+(\S+)\s+#/) {
	    $type  = $1;
	    $value = $2;
	    $name  = $3;

	    my $newvalue = $value;
	    
	    $ref_motif_name->[$nmotif]  = $name;
	    $ref_motif_value->[$nmotif] = $newvalue;
	    $ref_motif_type->[$nmotif]  = $type;
	    $ref_motif_idx->{$name}     = $nmotif;
	    $nmotif    += 1;
	    $nmotif_IL += 1;
	}
	elsif (/^\s*(J3)\s+(\S+\s+\S+\s+\S+)\s+(\S+)\s*$/ || /^\s*(J3)\s+(\S+\s+\S+\s+\S+)\s+(\S+)\s+#/) {
	    $type  = $1;
	    $value = $2;
	    $name  = $3;

	    my $newvalue = $value;
	    
	    $ref_motif_name->[$nmotif]  = $name;
	    $ref_motif_value->[$nmotif] = $newvalue;
	    $ref_motif_type->[$nmotif]  = $type;
	    $ref_motif_idx->{$name}     = $nmotif;
	    $nmotif    += 1;
	    $nmotif_J3 += 1;
	}
	elsif (/^\s*(J4)\s+(\S+\s+\S+\s+\S+\s+\S+)\s+(\S+)\s*$/ || /^\s*(J4)\s+(\S+\s+\S+\s+\S+\s+\S+)\s+(\S+)\s+#/) {
	    $type  = $1;
	    $value = $2;
	    $name  = $3;

	    my $newvalue = $value;
	    
	    $ref_motif_name->[$nmotif]  = $name;
	    $ref_motif_value->[$nmotif] = $newvalue;
	    $ref_motif_type->[$nmotif]  = $type;
	    $ref_motif_idx->{$name}     = $nmotif;
	    $nmotif    += 1;
	    $nmotif_J4 += 1;
	}
	elsif (/^\s*(BS)\s+(\S+)\s+(\S+)\s*$/ || /^\s*(BS)\s+(\S+)\s+(\S+)\s+#/) {
	    $type  = $1;
	    $value = $2;
	    $name  = $3;

	    my $newvalue = $value;
	    
	    $ref_motif_name->[$nmotif]  = $name;
	    $ref_motif_value->[$nmotif] = $newvalue;
	    $ref_motif_type->[$nmotif]  = $type;
	    $ref_motif_idx->{$name}     = $nmotif;
	    $nmotif    += 1;
	    $nmotif_BS += 1;
	    
	    $name .= ".rev";
	    $ref_motif_name->[$nmotif]  = $name;
	    $ref_motif_value->[$nmotif] = $newvalue;
	    $ref_motif_type->[$nmotif]  = $type;
	    $ref_motif_idx->{$name}     = $nmotif;
	    $nmotif += 1;
	    $nmotif_BS += 1;
	}
    }
    close(FILE);

    $$ret_nmotif    = $nmotif;
    $$ret_nmotif_HL = $nmotif_HL;
    $$ret_nmotif_BL = $nmotif_BL;
    $$ret_nmotif_IL = $nmotif_IL;
    $$ret_nmotif_J3 = $nmotif_J3;
    $$ret_nmotif_J4 = $nmotif_J4;
    $$ret_nmotif_BS = $nmotif_BS;
}

sub motifs_table {
    my ($file, $nmotif_HL, $nmotif_BL, $nmotif_IL, $nmotif_J3, $nmotif_J4, $nmotif_BS,
	$nmotif, $ref_motif_name, $ref_motif_value, $ref_motif_type, $ref_motif_idx) = @_;

    open (TBL, ">$file") || die;

    my $HL_header = "\\textbf\{\\#HL\} &  \\textbf\{Loop(5'-3')\} & \\textbf\{L(5'-3')\} & \\textbf\{R(5'-3')\} &&&& \\textbf\{\} \\\\";
    my $BL_header = "\\textbf\{\\#BL\} &  \\textbf\{Loop(5'-3')\} & \\textbf\{L(5'-3')\} & \\textbf\{R(5'-3')\} &&&& \\textbf\{\} \\\\";
    my $IL_header = "\\textbf\{\\#IL\} &  \\textbf\{Loop-L(5'-3')\} & \\textbf\{Loop-R(5'-3')\} & \\textbf\{L-o(5'-3')\} & \\textbf\{R-o(5'-3')\} & \\textbf\{L-i(5'-3')\} & \\textbf\{R-i(5'-3')\} & \\textbf\{\}\\\\";
    my $J3_header = "\\textbf\{\\#J3\} &  \\textbf\{S1(5'-3')\} & \\textbf\{S2(5'-3')\} & \\textbf\{S3(5'-3')\} &&&& \\textbf\{\}\\\\";
    my $J4_header = "\\textbf\{\\#J4\} &  \\textbf\{S1(5'-3')\} & \\textbf\{S2(5'-3')\} & \\textbf\{S3(5'-3')\} & \\textbf\{S4(5'-3')\} &&& \\textbf\{\}\\\\";
    my $BS_header = "\\textbf\{\\#BS\} &  \\textbf\{S(5'-3')\} &&&&&& \\textbf\{\}\\\\";

    print TBL "\\begin\{longtable\}\{|llllllll|\}\n";
    print TBL "\\caption*\{\\footnotesize Table 3. \\texttt\{\\bf Motifs in CaCoFold-R3D.\} The descriptor file includes 56 different motif architectures, and a total of 96 variants. HL = Hairpin Loop, BL = Bulge Loop, IL=Internal Loop, J3 = 3-way Junction, J4 = 4-way Junction, BS = Branch Segment.\}\\\\";
    print TBL "\\hline\n \\textbf\{Motif Type\} & \\multicolumn\{6\}\{c\}\{\\textbf\{Motif descriptor\}\} & \\textbf\{Motif Name\}\\\\\n"; 
    print TBL "\\hline\n$HL_header\n\\hline\n"; 
    for (my $m = 0; $m < $nmotif; $m ++) {
	if    ($m == $nmotif_HL) { print TBL "\\hline\n$BL_header\n\\hline\n"; }
	elsif ($m == $nmotif_HL+$nmotif_BL) { print TBL "\\hline\n$IL_header\n\\hline\n"; }
	elsif ($m == $nmotif_HL+$nmotif_BL+$nmotif_IL) { print TBL "\\hline\n$J3_header\n\\hline\n"; }
	elsif ($m == $nmotif_HL+$nmotif_BL+$nmotif_IL+$nmotif_J3) { print TBL "\\hline\n$J4_header\n\\hline\n"; }
	elsif ($m == $nmotif_HL+$nmotif_BL+$nmotif_IL+$nmotif_J3+$nmotif_J4) { print TBL "\\hline\n$BS_header\n\\hline\n"; }

	my $type  = $ref_motif_type->[$m];
	my $name  = $ref_motif_name->[$m]; $name =~ s/\_/\\\_/g; $name =~ s/\-/\\\_/g;$name =~ s/\^/\\\^/g;
	my $value = $ref_motif_value->[$m];

	my @value = split ' ', $value;
	my $len = $#value+1;
	my $newvalue = "";
	for (my $n = 0; $n < 6; $n ++) {
	    if ($n < $len) { my $val = $value[$n]; $newvalue .= "$val & "; }
	    else           { $newvalue .= " & "; }
	}
	print TBL "$type & $newvalue  $name\\\\\n";
    }
    print TBL "\\hline\n";
    print TBL "\\end\{longtable\}\n";
   
    close(TBL);
}
```
