## Supplementary material for "All-at-once RNA folding with 3D motif prediction framed by evolutionary information": R-scape_userguide.pdf

### R-scape User's Guide

RNA Significant Covariation Above Phylogenetic Expectation

Version v2.5.7; March 2025

Elena Rivas

Department of Molecular and Celullar Biology
Harvard University
52 Oxford Street, Northwest Building #430
Cambridge MA 02138 USA
http://rivaslab.org/

Copyright (C) 2017-2025 Elena Rivas, Harvard University.

Permission is granted to make and distribute verbatim copies of this manual provided the copyright notice and this permission notice are retained on all copies.

R-scape is licensed and freely distributed under the GNU General Public License version 3 (GPLv3). For a copy of the License, see `http://www.gnu.org/licenses/`.

**10 Acknowledgments** **47**

### 1 Introduction

R-scape (RNA Significant Covariation Above Phylogenetic Expectation) is a program that given a multiple sequence alignment (MSA) of RNA sequences, finds the pairs of positions that show a pattern of significant covariation. Each covariation score has an E-value associated to it. E-values are determined using a null model of covariation due to phylogeny but independent of any structural constraints.

#### How to avoid reading this manual

- Follow the quick installation instructions on page 6.
- Go to the tutorial section on page 8, which walks you through some examples of using R-scape on real data.

Everything else, you can read later.

#### How do I cite R-scape?

Rivas, E. *et al.*, *"A statistical test for conserved RNA structure shows lack of evidence for structure in lncRNAs"*, Nature Methods 14, 4548 (2017).
`https://www.nature.com/articles/nmeth.4066`
Rivas, E. and Eddy, S. E., *"Response to Tavares et al., Covariation analysis with improved parameters reveals conservation in lncRNA structures"*, (2018).
`https://doi.org/10.1101/2020.02.18.955047`.
Rivas, E. *et al.*, *"Estimating the power of sequence covariation analysis for detecting conserved RNA structure"*, *Bionformatics*, 36, 30723076, (2020).
`https://doi.org/10.1093/bioinformatics/btaa080`
Rivas, E., *"RNA covariation at helix-level resolution for the identification of evolutionarily conserved RNA structure"*, *PLOS Comput Biol, 19(7): e1011262, (2023).*
`https://doi.org/10.1371/journal.pcbi.1011262`.

#### How do I cite CaCoFold?

Rivas, E., *"RNA structure prediction using positive and negative evolutionary information"*, *PLOS Comput Biol*, 16(10), e1008387, (2020).
`https://doi.org/10.1371/journal.pcbi.1008387`.

#### How do I cite CaCoFold-R3D?

Karan, A and Rivas, E., *"All-at-once RNA folding with 3D motif prediction framed by evolutionary information"*, *bioRxiv*, Dec (2024).
`https://doi.org/10.1101/2024.12.17.628809`, 20 Dec 2024.

### 2 Installation

#### Quick installation instructions

Download `R-scape.tar.gz` from `http://eddylab.org/`; unpack it, configure, and make:

```
> tar xf R-scape.tar.gz
> cd R-scape
> ./configure
> make
> make install
```

The newly compiled binary (`R-scape`) is in the `R-scape/bin` directory. You can run it from there, as in this example:

```
> bin/R-scape tutorial/updated_Arisong.sto
```

That's it. You can keep reading if you want to know more about customizing a R-scape installation, or you can skip ahead to the next chapter, the tutorial.

#### System requirements

**Operating system:** R-scape is designed to run on POSIX-compatible platforms, including UNIX, Linux and Mac OS/X. The POSIX standard essentially includes all operating systems except Microsoft Windows. We have tested most extensively on Linux and MacOS/X because these are the machines we develop on.

**Compiler:** The source code is C conforming to POSIX and ANSI C99 standards. It should compile with any ANSI C99 compliant compiler, including the GNU C compiler `gcc`, and the C++ compiler `g++`. We test the code using the `gcc` and `g++` compilers.

The code include several Perl scripts (from the independent program R2R used here). Make sure your PATH environmental variable includes a directory with a Perl executable.

The code also uses GNUPLOT. Make sure your PATH environmental variable includes a directory with a GNUPLOT executable.

**Libraries and other installation requirements:** R-scape includes three software libraries:

- the Easel library package (`http://bioeasel.org/`),
- the HMMER library package (`http://hmmer.org/`),
- the Infernal library package (`http://eddylab.org/infernal/`),

and three independent programs:

- FastTree (Price et al., 2010) (for building phylogenetic trees, v2.1.11),
- R2R (Weinberg and Breaker, 2011) (for drawing consensus RNA structures),
- RNAVIEW (Yang et al., 2003) (for identifying different types of basepairs in nucleic acid alignments).

All libraries and independent programs will automatically compile during R-scape's installation process. By default, R-scape does not require any additional libraries to be installed by you, other than standard ANSI C99 libraries that should already be present on a system that can compile C code.

Executables for the three independent programs will appear in the `R-scape/bin` directory.

#### Makefile targets

**all** Builds everything. Same as just saying `make`.

**install** Installs the binaries (`R-scape`, `FastTree`, `r2r`).

By default, programs are installed in `R-scape_version/bin`. You can customize the location of the binaries by replacing

**> ./configure**

with

**> ./configure --prefix=/the/directory/you/want**

The newly compiled binaries are now in the `/the/directory/you/want/bin` directory.

**uninstall** Reverses the steps of make install.

**clean** Removes all files generated by compilation (by `make`). Configuration (files generated by `./configure`) is preserved.

**distclean** Removes all files generated by configuration (by `./configure`) and by compilation (by `make`).

#### Why is the output of 'make' so clean?

Because we're hiding what's really going on with the compilation with a wrapper. If you want to see what the command lines really look like, pass a `V=1` option (V for "verbose") to `make`, as in:

**> make V=1**

#### What gets installed by 'make install', and where?

The top-level configure file has a variable RSCAPE_HOME that specifies the directory where `make install` will install things: `RSCAPE_HOME/bin`.

By default RSCAPE_HOME is assigned to the current directory R-scape.

The best way to change this default is when you use `./configure`, and the most important variable to consider changing is `--prefix`. For example, if you want to install R-scape in a directory hierarchy all of its own, you might want to do something like:

**> ./configure --prefix=/usr/local/rscape**

That would keep R-scape out of your system-wide directories like `/usr/local/bin`, which might be desirable. Of course, if you do it that way, you'd also want to add `/usr/local/rscape/bin` to your `$PATH`.

### 3 Tutorial for running R-scape

Here's a tutorial walk-through of how to use R-scape. This should suffice to get you started.

#### Modes of R-scape

For an input alignment, R-scape reports all pairs that have covariation scores with E-values smaller than a target E-value.

R-scape has two different **modes** of operation which determine how it calculates E-values, for which it needs to know how many possible base pairs were tested (i.e. E-values are multiple-test-corrected). The E-values are calculated in one of two ways:

**A one-set statistical test:** *default*

E-values are calculated assuming that all pairs are possible.
This is the default behavior of R-scape.

**A two-set statistical test:** `option -s`

If the alignment has associated a *given structure*, **`option -s`** performs two independent statistical tests:
one for the pairs included in the structure, a different one for all the remaining possible pairs.
It also draws the given consensus structure annotated with the significantly covarying base pairs.

#### The four options to run R-scape

These are the four options to run R-scape.

| | |
|---|---|
| Evaluate region for conserved structure | All possible pairs are analyzed equally as a one set test. If a consensus structure is provided, that structure is ignored in the covariation test, but it is visualized with the significant covarying pairs highlighted in green.<br>**preferred use:**<br>This option is most appropriate if you're trying to determine if a conserved structure exists. |
| Predict new structure | All possible pairs are analyzed equally. A structure is predicted and visualized with the significant covarying pairs highlighted in green.<br>**preferred use:**<br>This option is most appropriate for obtaining a new consensus structure prediction based on covariation analysis. |
| Evaluate given structure | Requires that your Stockholm file has a proposed consensus structure annotation. Two independent covariation tests are performed, one on the set of proposed basepairs, the other on all other possible pairs. The given structure is visualized with significant covarying pairs highlighted in green.<br>**preferred use:**<br>This option is most appropriate for evaluating how well an independently proposed consensus structure is supported by covariation analysis. |
| Improve given structure | Requires that your Stockholm file has a proposed consensus structure annotation. Two independent covariation tests are performed, one on the set |

of proposed basepairs, the other on all other possible pairs. A new consensus structure is predicted and visualized with the significant covarying pairs highlighted in green.

**preferred use:**
This option is most appropriate for using covariation analysis to improve your current consensus structure.

I'll show examples of running each mode, using examples in the `tutorial/` subdirectory of the distribution.

#### Option –RAF(S) disallowed

The options to use the covariation measures RAF, RAFa, RAFp, RAFS, RAFSp, and RAFSa has been disallowed, unless they are used in combination with option –naive which reports the list of values for all possible pairs without any statistical significance associated to them.

The following disclamer appears otherwise.

**`> bin/R-scape --RAF tutorial/updated_Arisong.sto`**

```
DISCLAIMER: This measure can only be used in combination with the --naive option.

The --naive option reports a ranked list of scores for all possible
pairs without assigning E-values. RAF, RAFS and related measures
(RAFp, RAFa, RAFSp, RAFSa) cannot be used in combination with
R-scape's statistical test.

The RAF(S) statistics measure covariation and consistency. RAFS
assigns relatively high scores to pairs of alignment columns that are
consistent with base pairing even if there is no covariation at
all. The RAFS statistic was developed for the purpose of predicting
consensus RNA structures from alignments of sequences already presumed
to have a structure (Hofacker et al., 2002; Lindgreen et al.,
2006). For this purpose, both covariation and consistency are useful
cues. Distinguishing a conserved RNA structure from a conserved
primary sequence is a different problem that requires using a
statistic that does not systematically detect significant signals on
conserved primary sequence alone. That is R-scape's statistical
test. The R-scape statistical test can only be used with measures that
estimate covariation alone such as mutual information (MI) or G-test
(GT).
```

#### Files used in the tutorial

The subdirectory `/tutorial` in the R-scape distribution contains the files used in the tutorial.

The tutorial provides several examples of RNA structural alignments, all in Stockholm format:

**`updated_Arisong.sto`** Structural alignment of the ciliate Arisong RNA. This alignment is an updated version of the one published in (Jung et al., 2011).

**`ar14.sto`** Structural alignment of the $\alpha$-proteobacteria ncRNA ar14. This alignment is an updated version of the one published in (del Val et al., 2012).

**`manA.sto`** Alignment of the manA RNA motif (**?**Weinberg et al., 2010) provided in the Zasha Weinberg database (ZWD) (Weinberg, 2018).

**`RF00005.sto`** Rfam v12.0 (Nawrocki et al., 2015) seed alignment of tRNA.

**`RF00001-noss.sto`** Rfam v12.0 seed alignment of 5S rRNA, after removing the consensus secondary structure.

#### Running R-scape on one alignment file

To run R-scape with default parameters on alignment file `tutorial/updated_Arisong.sto` use:

**`> bin/R-scape tutorial/updated_Arisong.sto`**

The output is a list of the significantly covarying positions under the one-set test

```
# R-scape :: RNA Structural Covariation Above Phylogenetic Expectation
# R-scape 1.4.0 (Oct 2019)
# Copyright (C) 2016 Howard Hughes Medical Institute.
# Freely distributed under the GNU General Public License (GPLv3).
#-------------------------------------------------------------------------------------------------------
# MSA updated_Arisong_1 nseq 95 (95) alen 66 (150) avgid 65.82 (64.97) nbpairs 20 (20)
# One-set statistical test (all pairs are tested as equivalent)
#
#
# Method Target_E-val [cov_min,cov_max] [FP | TP True Found | Sen PPV F]
# GTp    0.05         [-9.78,121.66]     [0 | 2 20 2 | 10.00 100.00 18.18]
#
#       left_pos       right_pos        score          E-value       substitutions      power
#-------------------------------------------------------------------------------------------------------
*              98             106       121.65645        0.00241628        45               0.48
*             122             137       91.44593        0.038356        57               0.58
```

A star "*" in the first column indicates that the pair is part of the annotated structure in the `updated_Arisong.sto` file. A blank indicates a pair that is not compatible with the structure. A "∼" indicates an interaction not in the annotated structure but compatible with it (none in this example).

The Arisong RNA in `tutorial/updated_Arisong.sto` has a proposed secondary structure. Instead of testing all pairs as equivalent, we may want to test the significance of the given structure as a one set of pairs, and independently that of the rest of all possible pairs. In order to do a two-set test use:

**`> bin/R-scape -s tutorial/updated_Arisong.sto`**

The output is a list of the significantly covarying positions under the two-set test.

```
# R-scape :: RNA Structural Covariation Above Phylogenetic Expectation
# R-scape 1.4.0 (Oct 2019)
# Copyright (C) 2016 Howard Hughes Medical Institute.
# Freely distributed under the GNU General Public License (GPLv3).
#-------------------------------------------------------------------------------------------------------
# MSA updated_Arisong_1 nseq 95 (95) alen 66 (150) avgid 65.82 (64.97) nbpairs 20 (20)
# Two-set statistical test (one test for annotated basepairs, another for all other pairs)
#
#
# Method Target_E-val [cov_min,cov_max] [FP | TP True Found | Sen PPV F]
# GTp    0.05         [-9.78,121.66]     [0 | 11 20 11 | 55.00 100.00 70.97]
#
#       left_pos       right_pos        score          E-value       substitutions      power
#-------------------------------------------------------------------------------------------------------
*              98             106       121.65645        2.25295e-05        45               0.48
*             122             137       91.44593        0.000357632        57               0.58
*              96             108       88.43400        0.000466924        26               0.28
*             120             139       74.80289        0.00162024        87               0.76
*             119             140       58.72158        0.00678565        90               0.78
*             121             138       58.34837        0.00691674        99               0.82
*              94             110       57.27959        0.00760538        37               0.40
*             124             134       55.67692        0.0086606        20               0.21
*             123             135       54.59630        0.00946822        72               0.68
*              99             105       53.44797        0.0107226        15               0.14
*              97             107       44.91842        0.0405594        58               0.59

# The given structure
# SS_cons ::::::::::::::::::::::::::::::::::::::::::::::::::::::::::::
#
# SS_cons :::::::::::::::::::::::::::::::::<<<<<<<___>>>>>>><<<<<<--<<
#
# SS_cons <<<<<_______>>>->>>>-->>>>>>::
#
```

```
# Power analysis of given structure
#
# covary  left_pos      right_pos    substitutions      power
#----------------------------------------------------------------
     *    94              110              37               0.40
          95              109              28               0.31
     *    96              108              26               0.28
     *    97              107              58               0.59
     *    98              106              45               0.48
     *    99              105              15               0.14
          100              104              20               0.21
          111              148              0               0.00
          112              147              18               0.18
          113              146              1               0.00
          114              145              15               0.14
          115              144              49               0.52
          116              143              106               0.84
     *    119              140              90               0.78
     *    120              139              87               0.76
     *    121              138              99               0.82
     *    122              137              57               0.58
     *    123              135              72               0.68
     *    124              134              20               0.21
          125              133              31               0.34
#
# BPAIRS 20
# avg substitutions per BP  43.7
# BPAIRS expected to covary 8.3
# BPAIRS observed to covary 11
```

The scores of the pairs are identical to those in the one-set test. The E-values have changed relative to those of the one-set test.

#### Default parameters

Default parameters are:

**Target E-value:** default is 0.05. R-scape reports pairs which covariation score has E-value smaller or equal to the target value. The target E-value can be changed with option **-E <x>**, $x >= 0$.

**Sequence weighting:** Sequences are weighted according to the Gerstein/Sonnhammer/Chothia (GSC) algorithm (Gerstein et al., 1994). This algorithm is time consuming. For alignments with more than 1000 sequences, we use the faster position-based weighting algorithm (Henikoff and Henikoff, 1994). Both weighting algorithms are implemented as part of the easel library.

**Gaps in columns:** Columns with more than 50% gaps are removed. The gap threshold for removing columns can be modified using option **--gapthresh <x>** , $0 < x <= 1$.

**Covariation statistic:** The default covariation statistic is the average product corrected G-Test (equivalent to option **--GTp**).

**Covariation Class:** R-scape uses the 16 component covariation statistic (C16), unless the number of sequences in the alignment is $\leq 8$ or the length of the alignment is $\leq 50$, in which case it uses the two-class covariation statistic (C2). A particular covariation class can be selected using either **--C16** or **--C2**.

The threshold for the minimum number of sequences can be changed with option `--nseqthresh <n>`. The threshold for the minimum alignment length can be changed with option `--alenthresh <n>`.

**`Null alignments:`** In order to estimate E-values, R-scape produces 20 null alignments, unless the product of the number of sequences by the length of the alignment $< 10{,}000$ in which case the number of null alignments is 50; or $< 1{,}000$ in which case it is 100. The number of null alignments can be controlled with option **`--nshuffle <n>`**.

A full list of the R-scape options is found by using

```
> R-scape -h
```

### 4 Tutorial for running CaCoFold

Here's a tutorial walk-through of how to use CaCoFold. CaCoFold uses the evolutionary information capture by R-scape in the form of the base pairs that significantly covary, and builds a structure. The CaCoFold structure may contain pseudoknots and triplets, provided that they present covariation signal.

#### The –cacofold option

After performing one of the two statistical tests with R-scape, this option implements the CaCoFold algorithm:

Builds the best consensus structure that includes the largest possible number of significantly covarying pairs, *the maximum-covariation optimal consensus structure*. The algorithm identifies pseudoknots and other not nested interactions by running a cascade of nested algorithms until all covarying pairs are taken into account.

Draws the *maximum-covariation optimal consensus structure* annotated with the significantly covarying base pairs.

It also returns the alignment in Stockholm format annotated with the max-cov optimal consensus structure.

```
> bin/R-scape --cacofold tutorial/updated_Arisong.sto
```

The output includes first the same output as default R-scape alone, followed by R-scape's proposed structure that under the heading "# The predicted CaCoFold structure" as follows,

```
# The predicted CaCoFold structure
# SS_cons ::::::::::::::::::::::::::::::::::::::::::::::::::::::::::::
#
# SS_cons ::::::::::::::::::::::::::::::::::<<<<<_____>>>>>:<<<<<---<<
#
# SS_cons <<<<_________>>->>>>--->>>>>::
#

# Power analysis of CaCoFold structure
#
# covary  left_pos      right_pos    substitutions      power
#-------------------------------------------------------------------
          95             109              28               0.31
          96             108              26               0.28
          97             107              58               0.59
     *    98             106              45               0.48
          99             105              15               0.14
          111             148              0               0.00
          112             147              18               0.18
          113             146              1               0.00
          114             145              15               0.14
          115             144              49               0.52
          119             140              90               0.78
          120             139              87               0.76
          121             138              99               0.82
     *    122             137              57               0.58
          123             135              72               0.68
          124             134              20               0.21
#
# BPAIRS 16
# avg substitutions per BP  42.5
# BPAIRS expected to covary 6.5
# BPAIRS observed to covary 2
```

The structure predicted by R-scape includes all the basepairs reported as covarying, provided that those can be arranged into one single structure (including pseudoknots and other non Watson-Crick interactions). The R-scape folding algorithm cannot deal with residues that covary with more than one other residue, such as is the case for alternative structures or triplets.

Similarly using

```
> bin/R-scape -s --cacofold tutorial/updated_Arisong.sto
```

The output includes first the same output as **`option -s`** of R-scape alone, followed by R-scape's proposed CaCoFild structure including all the the covarying pairs obtained under the two-set test.

```
# R-scape :: RNA Structural Covariation Above Phylogenetic Expectation
# R-scape 1.4.0 (Oct 2019)
# Copyright (C) 2016 Howard Hughes Medical Institute.
# Freely distributed under the GNU General Public License (GPLv3).
#-------------------------------------------------------------------------------------------------------
# MSA updated_Arisong_1 nseq 95 (95) alen 66 (150) avgid 65.82 (64.97) nbpairs 20 (20)
# Two-set statistical test (one test for annotated basepairs, another for all other pairs)
#
#
# Method Target_E-val [cov_min,cov_max] [FP | TP True Found | Sen PPV F]
# GTp    0.05         [-9.78,121.66]     [0 | 11 20 11 | 55.00 100.00 70.97]
#
#       left_pos       right_pos        score          E-value       substitutions      power
#-------------------------------------------------------------------------------------------------------
*            98            106       121.65645        2.25295e-05        45                0.48
*           122            137       91.44593        0.000357632        57                0.58
*            96            108       88.43400        0.000466924        26                0.28
*           120            139       74.80289        0.00162024        87                0.76
*           119            140       58.72158        0.00678565        90                0.78
*           121            138       58.34837        0.00691674        99                0.82
*            94            110       57.27959        0.00760538        37                0.40
*           124            134       55.67692        0.0086606        20                0.21
*           123            135       54.59630        0.00946822        72                0.68
*            99            105       53.44797        0.0107226        15                0.14
*            97            107       44.91842        0.0405594        58                0.59

# The given structure
# SS_cons ::::::::::::::::::::::::::::::::::::::::::::::::::::::::::::
#
# SS_cons ::::::::::::::::::::::::::::::::<<<<<<<___>>>>>>><<<<<<--<<
#
# SS_cons <<<<<_______>>>->>>>-->>>>>>::
#

# Power analysis of given structure
#
# covary  left_pos      right_pos    substitutions      power
#----------------------------------------------------------------
     *    94              110              37              0.40
          95              109              28              0.31
     *    96              108              26              0.28
     *    97              107              58              0.59
     *    98              106              45              0.48
     *    99              105              15              0.14
          100              104              20              0.21
          111              148              0              0.00
          112              147              18              0.18
          113              146              1              0.00
          114              145              15              0.14
          115              144              49              0.52
          116              143              106              0.84
     *    119              140              90              0.78
     *    120              139              87              0.76
     *    121              138              99              0.82
     *    122              137              57              0.58
     *    123              135              72              0.68
     *    124              134              20              0.21
          125              133              31              0.34
#
# BPAIRS 20
# avg substitutions per BP  43.7
# BPAIRS expected to covary 8.3
# BPAIRS observed to covary 11
#
#
# Method Target_E-val [cov_min,cov_max] [FP | TP True Found | Sen PPV F]
# GTp    0.05         [-9.78,121.66]     [0 | 11 17 11 | 64.71 100.00 78.57]
#
# in_fold in_given   left_pos       right_pos      score           E-value    substitutions      power
#---------------------------------------------------------------------------------------------------------------
*        *              98              106       121.65645        2.25295e-05        45                0.48
*        *             122              137       91.44593        0.000357632        57                0.58
*        *              96              108       88.43400        0.000466924        26                0.28
*        *             120              139       74.80289        0.00162024        87                0.76
```

```
*        *              119             140        58.72158       0.00678565       90               0.78
*        *              121             138        58.34837       0.00691674       99               0.82
*        *               94             110        57.27959       0.00760538       37               0.40
*        *              124             134        55.67692       0.0086606       20               0.21
*        *              123             135        54.59630       0.00946822       72               0.68
*        *               99             105        53.44797       0.0107226       15               0.14
*        *               97             107        44.91842       0.0405594       58               0.59

# The predicted CaCoFold structure
# SS_cons ::::::::::::::::::::::::::::::::::::::::::::::::::::::::::::
#
# SS_cons :::::::::::::::::::::::::::::::::<<<<<<_____>>>>>><<<<<---<<
#
# SS_cons <<<<_________>>->>>>--->>>>>::
#

# Power analysis of CaCoFold structure
#
# covary  left_pos      right_pos    substitutions      power
#----------------------------------------------------------------
     *    94              110              37              0.40
          95              109              28              0.31
     *    96              108              26              0.28
     *    97              107              58              0.59
     *    98              106              45              0.48
     *    99              105              15              0.14
          111              148              0              0.00
          112              147              18              0.18
          113              146              1              0.00
          114              145              15              0.14
          115              144              49              0.52
     *    119              140              90              0.78
     *    120              139              87              0.76
     *    121              138              99              0.82
     *    122              137              57              0.58
     *    123              135              72              0.68
     *    124              134              20              0.21
#
# BPAIRS 17
# avg substitutions per BP  42.2
# BPAIRS expected to covary 6.9
# BPAIRS observed to covary 11
```

#### Example of an RNA with pseudoknots

R-scape implements the CaCoFold folding algorithm capable of predicting pseudoknots and other non nested interactions using a cascade of dynamic programming algorithms. R-scape had adapted the program R2R to automatically include in the display all covarying interactions whether they are nested or not.

Consider the manA RNA motif. Both the proposed structure for manA RNA and the predicted CaCoFold structure have 2 pseudoknots with covariation support:

```
> bin/R-scape -s --cacofold tutorial/manA.sto
```

```
# The given structure
# SS_cons   <<<<<<_______________________________>>>>>>:::::[[[[[,,,<<--
# SS_cons_1 ::::::::::::::::::::::::::::::::::::::::::::::::::::::::::::
# SS_cons_2 ::::::::::::::::::::::::::::::::::::::::::::::::::::::::::::
#
# SS_cons   -----------<<<<<<__________________>>>>>>---->>,,,,((-((((,,
# SS_cons_1 ::::::::::::::::::::::::::::::::::::::::::::::::::::::::::::
# SS_cons_2 ::::::::::::::::::::::::::::::::::::::::::::::::::::::::::::
#
# SS_cons   ,,,,,,,,,,,,,,,,,,,,,,,,,,,,,<<<_______>>>,<<<<<<___________
# SS_cons_1 ::::::::::::::::::::::::::::::::::<<<<______________________
# SS_cons_2 ::::::::::::::::::::::::::::::::::::::::::::::::::::::::::::
#
# SS_cons   ____________________>>>>>>,,,<<<<<____________>>>>>,,))))--)
# SS_cons_1 _______________________________________>>>>:::::::::::::::::
# SS_cons_2 ::::::::::::::::::::::::::::::::::::::::::::::::::::::::::::
#
# SS_cons   ),,,,<<<<<<<------------<<<<____________________________>>>>
# SS_cons_1 ::::::::::::::::::::::::::::::::::::::::::::::::::::::::::::
```

```
# SS_cons_2 :::::::::::::::::::::::::::::::::<<<-<______________________
#
# SS_cons   ------->>>>>>>,,<<<<<_____________>>>>>]]]]]::::::
# SS_cons_1 ::::::::::::::::::::::::::::::::::::::::::::::::::
# SS_cons_2 _________________________>>>>:::::::::::::::::::::
#

# Power analysis of given structure
#
# covary  left_pos      right_pos    substitutions      power
#----------------------------------------------------------------
     *    1              43              61                0.61
     *    2              42              61                0.61
     *    3              41              72                0.68
     *    4              40              104                0.83
     *    5              39              106                0.84
     *    6              38              126                0.90
          49              344              15                0.14
     *    50              343              16                0.16
     *    51              342              26                0.28
     *    52              341              77                0.71
     *    53              340              26                0.28
     *    57              107              38                0.41
     *    58              106              16                0.16
     *    72              101              71                0.68
     *    73              100              38                0.41
     *    74              99              60                0.60
     *    75              98              34                0.37
     *    76              97              31                0.34
          77              96              27                0.30
     *    112              241              41                0.44
     *    113              240              51                0.53
     *    115              237              61                0.61
     *    116              236              46                0.49
     *    117              235              62                0.62
     *    118              234              49                0.52
     *    150              162              52                0.54
     *    151              161              47                0.50
     *    152              160              36                0.39
          155              223              31                0.34
     *    156              222              30                0.33
          157              221              28                0.31
          158              220              27                0.30
     *    164              206              29                0.32
     *    165              205              54                0.56
     *    166              204              148                0.94
     *    167              203              109                0.85
     *    168              202              154                0.95
     *    169              201              147                0.94
          210              231              48                0.51
     *    211              230              77                0.71
     *    212              229              71                0.68
     *    213              228              59                0.60
     *    214              227              75                0.70
     *    246              314              62                0.62
     *    247              313              92                0.79
     *    248              312              141                0.93
     *    249              311              90                0.78
     *    250              310              99                0.82
     *    251              309              105                0.84
     *    252              308              37                0.40
     *    265              300              61                0.61
     *    266              299              44                0.47
     *    267              298              42                0.45
     *    268              297              36                0.39
     *    274              329              47                0.50
     *    275              328              36                0.39
     *    276              327              39                0.42
          278              326              42                0.45
          317              339              13                0.12
     *    318              338              37                0.40
     *    319              337              60                0.60
     *    320              336              18                0.18
          321              335              0                0.00
#
# BPAIRS 63
# avg substitutions per BP  57.7
```

```
# BPAIRS expected to covary 33.2
# BPAIRS observed to covary 54

  ...

  ...

# The predicted CaCoFold structure
# SS_cons   <<<<<<_______________________________>>>>>>:[---[[[[[,,,<<--
# SS_cons_1 ::::::::::::::::::::::::::::::::::::::::::::::::::::::::::::
# SS_cons_2 ::::::::::::::::::::::::::::::::::::::::::::::::::::::::::::
#
# SS_cons   -----------<<<<<<__________________>>>>>>---->>,(--((-((((,,
# SS_cons_1 ::::::::::::::::::::::::::::::::::::::::::::::::::::::::::::
# SS_cons_2 ::::::::::::::::::::::::::::::::::::::::::::::::::::::::::::
#
# SS_cons   ,,,,,,,,,,,,,,,,,,,,,,,,,,,,,<<<_______>>>,<<<<<<___________
# SS_cons_1 ::::::::::::::::::::::::::::::::::<<<<______________________
# SS_cons_2 ::::::::::::::::::::::::::::::::::::::::::::::::::::::::::::
#
# SS_cons   ____________________>>>>>>,,,<<<<<---<____>--->>>>>,,))))--)
# SS_cons_1 _______________________________________>>>>:::::::::::::::::
# SS_cons_2 ::::::::::::::::::::::::::::::::::::::::::::::::::::::::::::
#
# SS_cons   )--),<<<<<<<------------<<<<-<<______>>----------------->>>>
# SS_cons_1 ::::::::::::::::::::::::::::::::::::::::::::::::::::::::::::
# SS_cons_2 :::::::::::::::::::::::::::::::::<<<-<______________________
#
# SS_cons   ------->>>>>>>,,<<<<<_____________>>>>>]]]]]-]::::
# SS_cons_1 ::::::::::::::::::::::::::::::::::::::::::::::::::
# SS_cons_2 _________________________>>>>:::::::::::::::::::::
#

# Power analysis of CaCoFold structure
#
# covary  left_pos      right_pos    substitutions      power
#-----------------------------------------------------------------
     *    1              43              61                0.61
     *    2              42              61                0.61
     *    3              41              72                0.68
     *    4              40              104                0.83
     *    5              39              106                0.84
     *    6              38              126                0.90
          45              346              82                0.74
          49              344              15                0.14
     *    50              343              16                0.16
     *    51              342              26                0.28
     *    52              341              77                0.71
     *    53              340              26                0.28
     *    57              107              38                0.41
     *    58              106              16                0.16
     *    72              101              71                0.68
     *    73              100              38                0.41
     *    74              99              60                0.60
     *    75              98              34                0.37
     *    76              97              31                0.34
          77              96              27                0.30
     *    109              244              37                0.40
     *    112              241              41                0.44
     *    113              240              51                0.53
     *    115              237              61                0.61
     *    116              236              46                0.49
     *    117              235              62                0.62
     *    118              234              49                0.52
     *    150              162              52                0.54
     *    151              161              47                0.50
     *    152              160              36                0.39
          155              223              31                0.34
     *    156              222              30                0.33
          157              221              28                0.31
          158              220              27                0.30
     *    164              206              29                0.32
     *    165              205              54                0.56
     *    166              204              148                0.94
     *    167              203              109                0.85
     *    168              202              154                0.95
     *    169              201              147                0.94
```

```
      210               231               48              0.51
   *  211               230               77              0.71
   *  212               229               71              0.68
   *  213               228               59              0.60
   *  214               227               75              0.70
      218               223               59              0.60
   *  246               314               62              0.62
   *  247               313               92              0.79
   *  248               312               141              0.93
   *  249               311               90              0.78
   *  250               310               99              0.82
   *  251               309               105              0.84
   *  252               308               37              0.40
   *  265               300               61              0.61
   *  266               299               44              0.47
   *  267               298               42              0.45
   *  268               297               36              0.39
      270               279               47              0.50
      271               278               81              0.73
   *  274               329               47              0.50
   *  275               328               36              0.39
   *  276               327               39              0.42
      278               326               42              0.45
      317               339               13              0.12
   *  318               338               37              0.40
   *  319               337               60              0.60
   *  320               336               18              0.18
      321               335               0              0.00
#
# BPAIRS 68
# avg substitutions per BP  58.0
# BPAIRS expected to covary 36.2
# BPAIRS observed to covary 55
```

The “SS_cons_1” and “SS_cons_2” lines describe the interactions that are not nested relative to the main “SS_cons” structure.

R-scape uses R2R to produce figures of the consensus structures where pseudoknots are also annotated. R-scape] option -s produces the file **tutorial/manA.R2R.sto.**{**pdf,svg**} with the structure annotated in the input alignment. R-scape] option –cacofold produces the file **tutorial/manA.fold.R2R.sto.**{**pdf,svg**} with the structure produced by R-scape. See Figure 1.

#### Single sequence structure prediction

If the alignment includes only one sequence, no statistical test is performed.

```
> bin/R-scape --cacofold tutorial/manA-oneseq.sto
```

reports the best structure given the sequence. No covariation support is possible for any of the basepairs reported from this analysis. Structures produced this way have to taken with great skepticism.

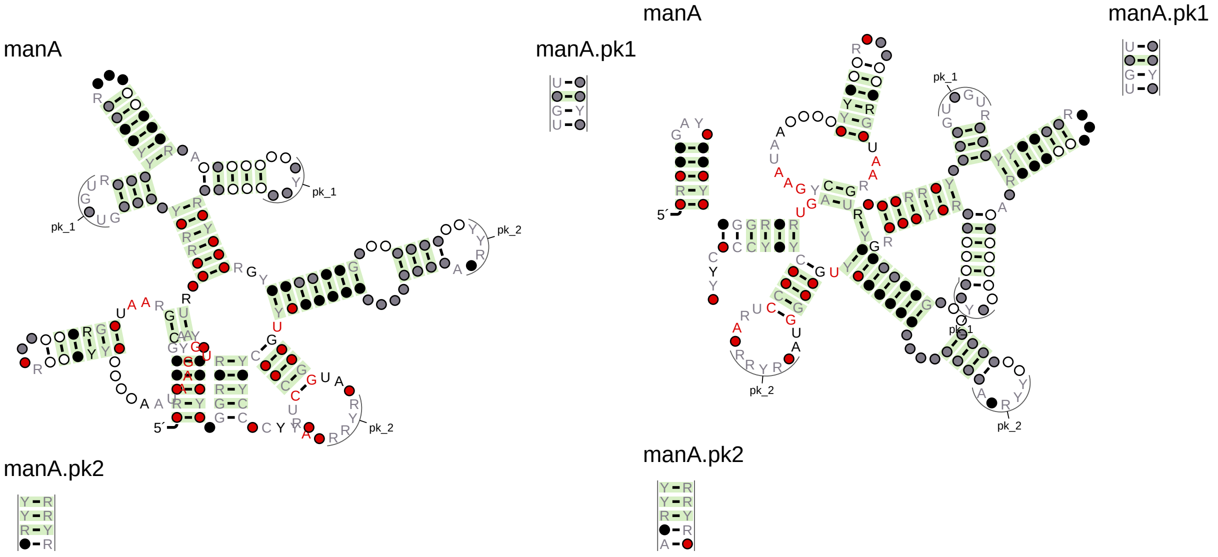

Figure 1: **Left: `tutorial/manA.R2R.sto.`{`pdf,svg`}**, the consensus secondary structure given in the input alignment, depicted by R-scape, using the program R2R. **Right: `tutorial/manA.fold.R2R.sto.`{`pdf,svg`}**, The consensus structure produced by R-scape (option –cacofold). Base pairs with covariation scores equal or below the target E-value (0.05 as default) are depicted in green.

### 5 Tutorial for running CaCoFold-R3D

Here's a tutorial walk-through of how to use CaCoFold-R3D. CaCoFold-R3D, extends CaCoFold by reporting a predicted structure that incorporates starndard Watson-Crick helices together with 3D RNA motifs that are composed of non-Watson-Crick pairs and unpaired residues. As with CaCoFold, the CaCoFold-R3D structure may contain pseudoknots and triplets, provided that they present covariation signal.

#### The –cacofold –r3d options

After performing one of the two statistical tests with R-scape, these two options together implements the CaCoFold-R3D algorithm.

#### Examples of 3D motifs and how to incorporate them to a R3D descriptor file

#### Hairpin Loops (HL)

Here we present the general HL SCFG model, and two example of HL mofits with their corresponding descriptor.

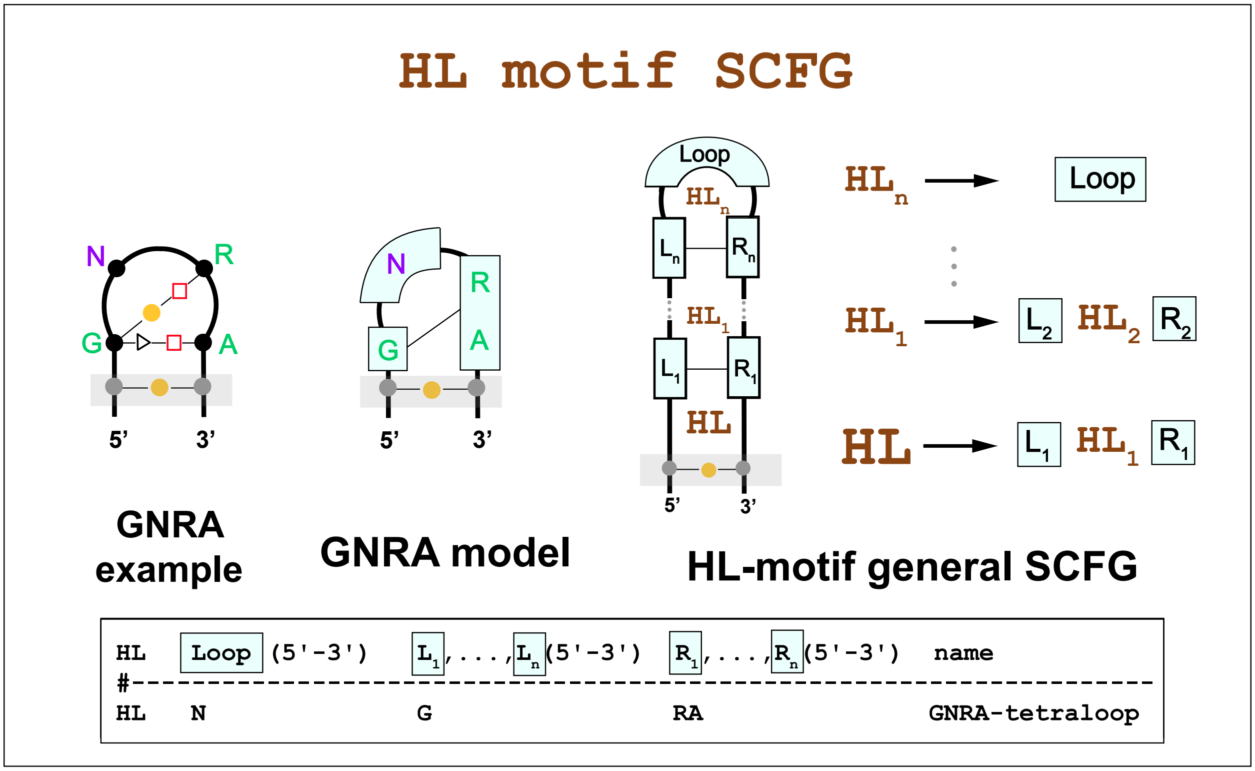

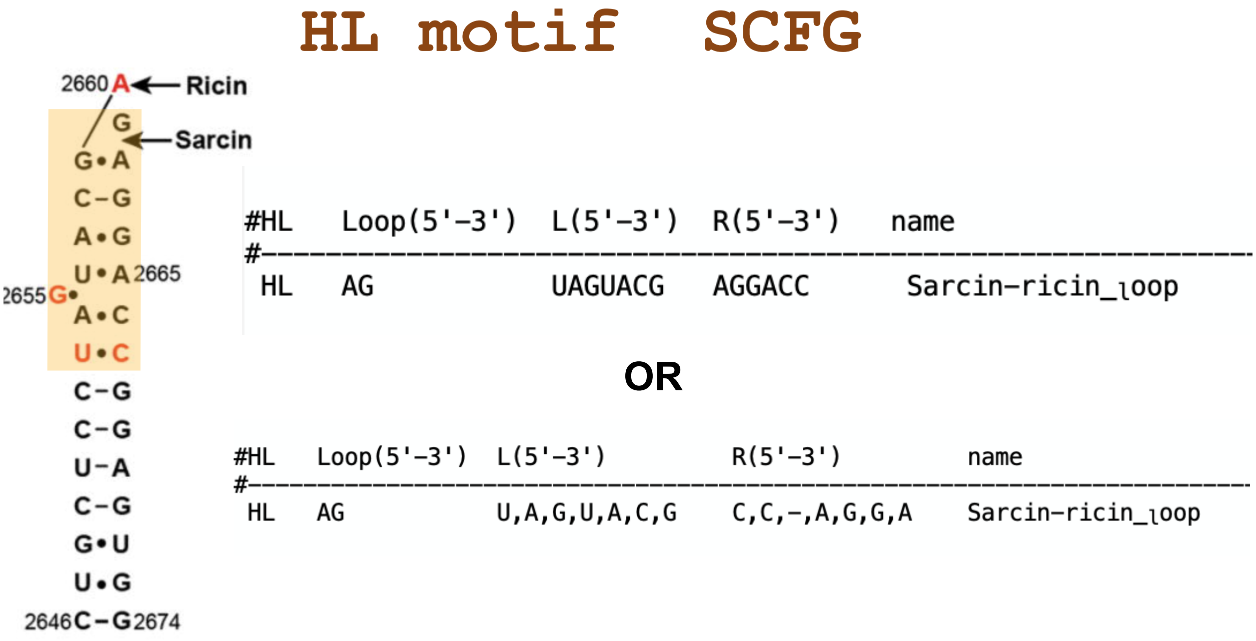

#### Bulge Loops (BL)

Here we present the general BL SCFG model, and two example of BL mofits with their corresponding descriptor.

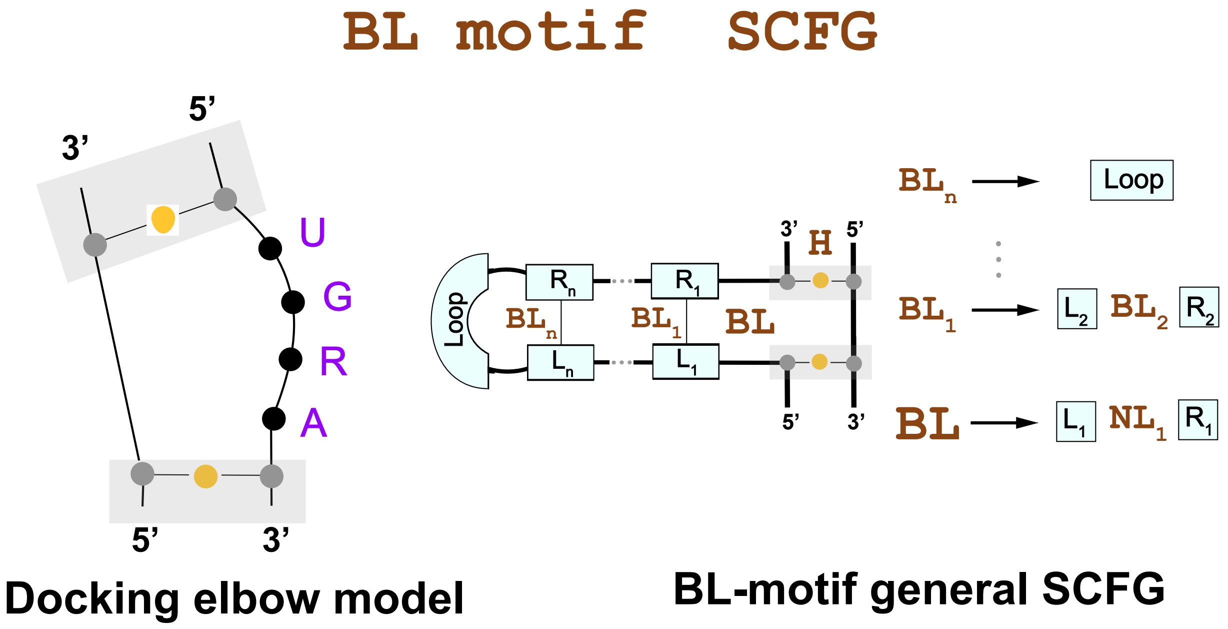

| #BL | Loop(5'-3') | L(5'-3') | R(5'-3') | name |
|---|---|---|---|---|
| BL | UGRAA | - | - | Docking-elbow |
| [illegible] | [illegible] | | | [illegible] |

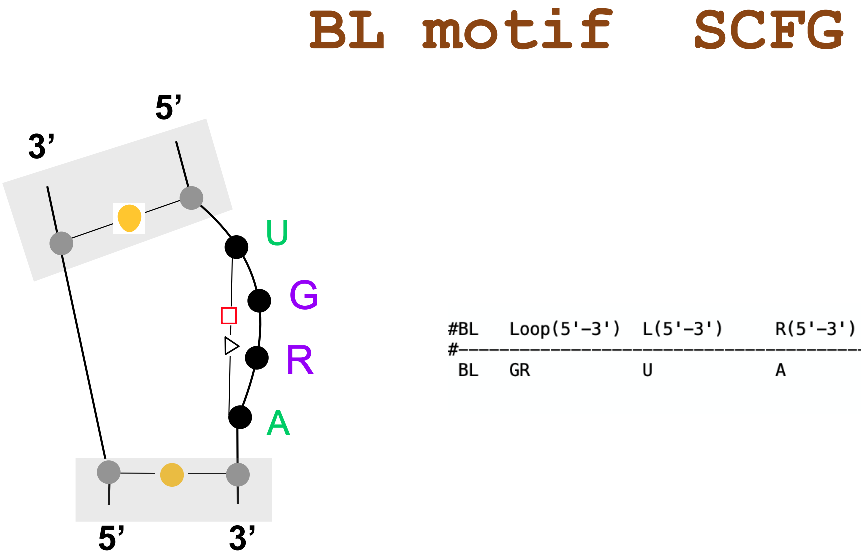

| #BL | Loop(5'-3') | L(5'-3') | R(5'-3') | name |
|---|---|---|---|---|
| BL | GR | U | A | Fake-bulge |

#### Internal Loops (IL)

Here we present the general IL SCFG model, and two example of IL mofits with their corresponding descriptor.

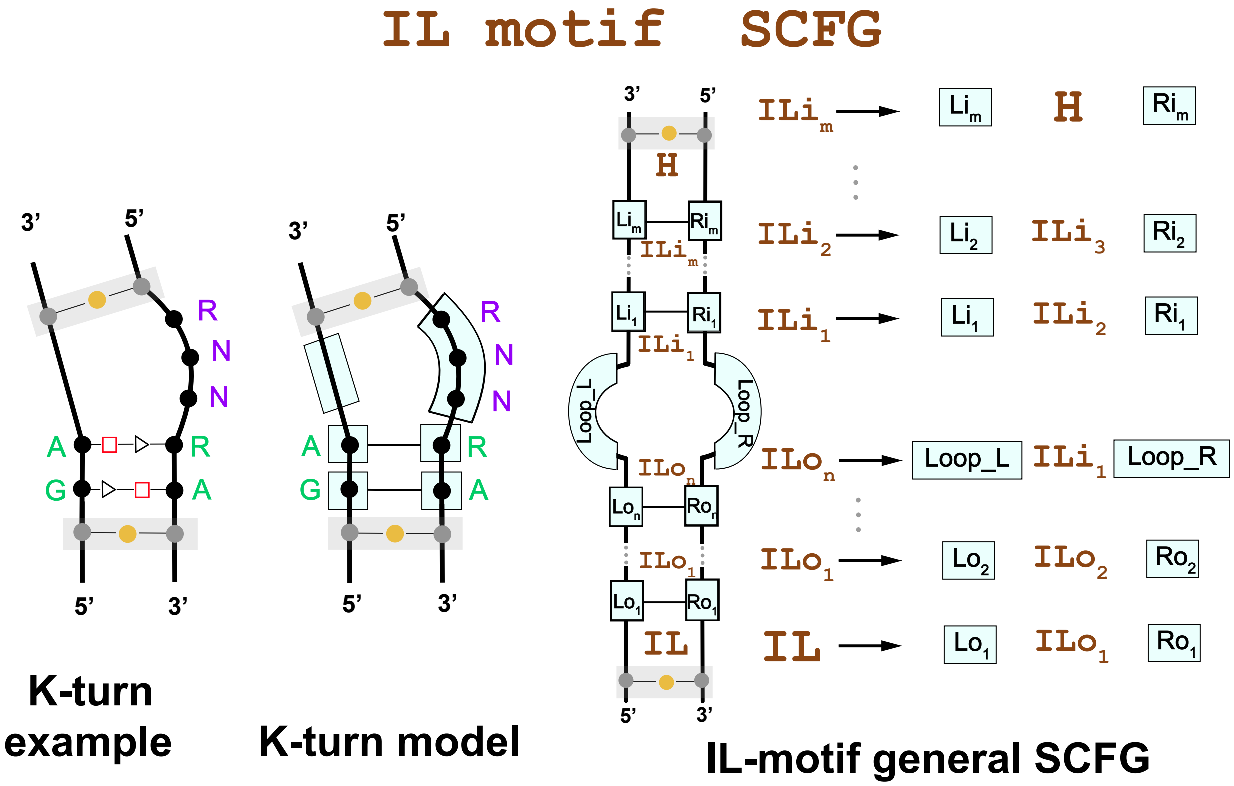

| #IL | Loop-L(5'-3') | Loop-R(5'-3') | L-o(5'-3') | R-o(5'-3') | L-i(5'-3') | R-i(5'-3') | name |
|---|---|---|---|---|---|---|---|
| IL | - | RNN | G,A | A,R | - | - | K-turn |

### IL motif SCFG

#### Loop-E

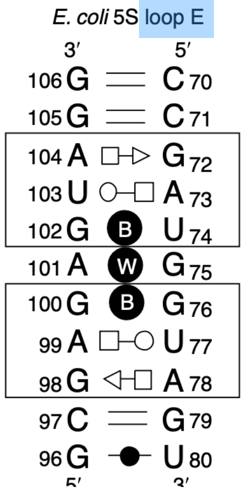

| #IL | Loop-L(5'-3') | Loop-R(5'-3') | L-o(5'-3') | R-o(5'-3') | L-i(5'-3') | R-i(5'-3') | name |
|---|---|---|---|---|---|---|---|
| IL | GAG | UGG | G,A | U,A | U,A | A,G | Loop-E |

#### hook-turn

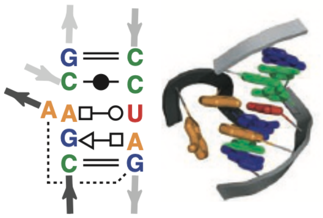

| #IL | Loop-L(5'-3') | Loop-R(5'-3') | L-o(5'-3') | R-o(5'-3') | L-i(5'-3') | R-i(5'-3') | name |
|---|---|---|---|---|---|---|---|
| IL | A | - | G,A | A,U | C | C | hook-turn |

##### 3-way Junctions (J3)

Here we present the general J3 SCFG model, and one example of J3 mofits with their corresponding descriptor.

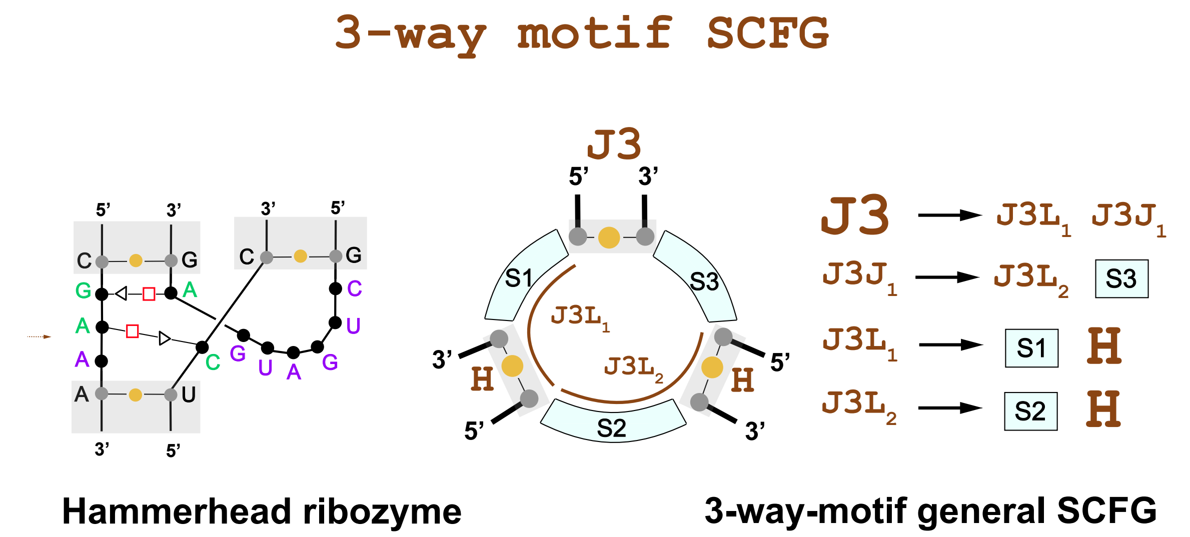

| J3 | S1(5'-3') | S2(5'-3') | S3(5'-3') | name |
|---|---|---|---|---|
| #---------------------------------------------------------------- |  |  |  |  |
| J3 | C | CUGAUGA | GAA | J3_hammerhead |

#### 4-way Junctions (J4)

Here we present the general J4 SCFG model, and one example of J4 mofits with their corresponding descriptor.
Example of a 4-way Junction.

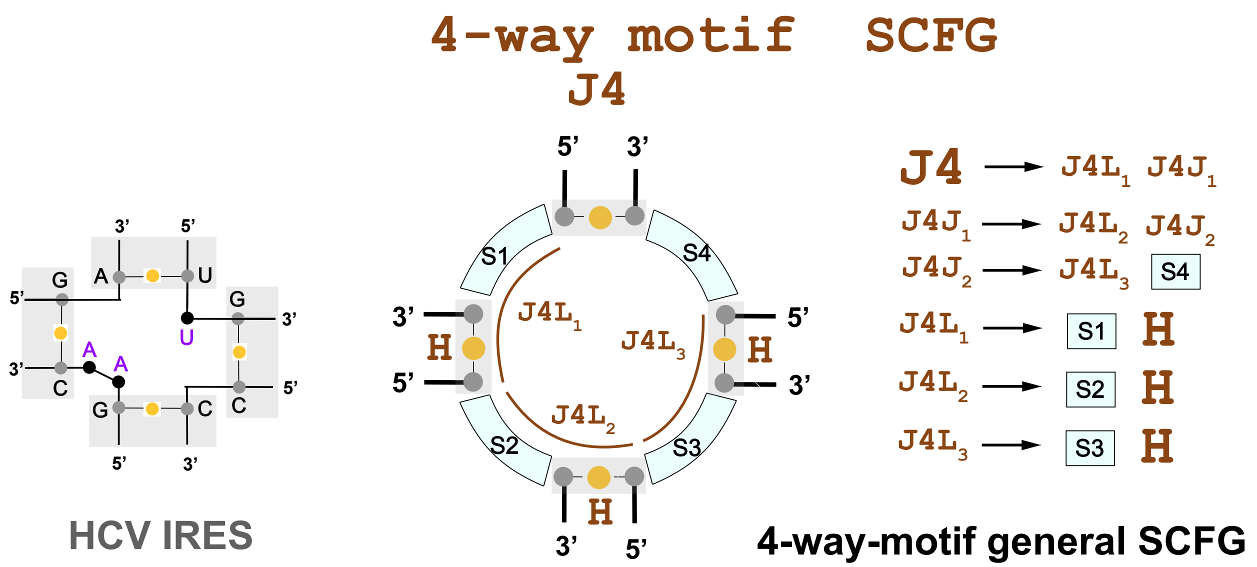

```
#J4    S1(5'-3')    S2(5'-3')   S3(5'-3')    S4(5'-3')       name
#----------------------------------------------------------------------
J4       -           AA          -            U               J4_HCV_IRES
```

##### Branch Segments (BS)

BS sequence segments can appear in any higher order junction. Here we provide the general form of a BS SCFG, and the particular case of a T-loop segnment.

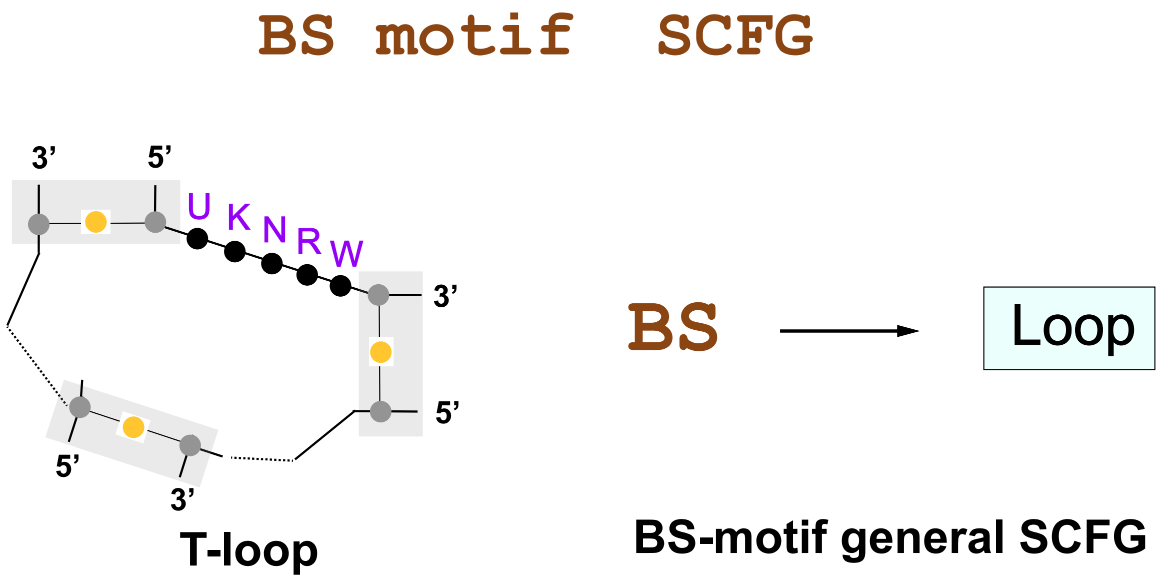

```
#BS   Loop(5'-3')   name
#-------------------------------------
BS        UKNRW              T-loop
```

#### The –r3dfile to customize the descriptor file

CaCoFold-R3D reads a descriptor file to identify all the 3D motifs, and to build the probabilistic model incorporating all of them. The default descriptor file used is given below (lines starting with # are comments).

Using the option `--r3dfile`, one can introduce any customized combination of 3D motifs to consider.

```
#HL   Loop(5'-3')  L(5'-3')  R(5'-3')    name
#------------------------------------------------------------------------------------------------------------------
 HL   N            G         RA          GNRA-tetraloop
 HL   URA          -         -           U-turn
 HL   UNCG         -         -           UNCG-tetraloop
 HL   ANYA         -         -           ANYA-tetraloop
 HL   CUYG         -         -           CUYG-tetraloop
 HL   YGNN         -         -           YGNN-tetraloop
 HL   GANC         -         -           GANC-tetraloop
 HL   UNAC         -         -           UNAC-tetraloop
 HL   URRR         -         -           T-loop-tetraloop
 HL   UAACR        -         -           L8_RNaseP_bact_a
 HL   AG           UAGUACG   AGACC       Sarcin-ricin_loop
 HL   AGGAY        -         -           CsrA_binding
 HL   GAGUA        -         -           GAGUA_pentaloop
 HL   AA           -         -           AA-5SrRNA
 HL   GCRYA        -         -           U6-loop

#BL   Loop(5'-3')  L(5'-3')  R(5'-3')   name
#------------------------------------------------------------------------------------------------------------------
 BL   UGRAA        -         -          Docking-elbow
 BL   AA           -         -          AA-bulge
 BL   G            -         -          bulged-G

#IL   Loop-L(5'-3') Loop-R(5'-3')  L-o(5'-3')     R-o(5'-3')     L-i(5'-3')   R-i(5'-3')     name
#------------------------------------------------------------------------------------------------------------------
 IL   RNN           -             G,A            A,R            -            -              K-turn
 IL   -             RNN           G,A            A,R            -            -              K-turn-b
 IL   N             -             R,A,R,R        A,U,R,R        -            -              Loop-E
 IL   UAAG          UAU           C,C            G,G            -            -              GAAA_Tetraloop-receptor
 IL   CVC           V             -              -              -            -              C-loop
 IL   G             -             G,A            A,A            U,A          A,G            G-bulge
 IL   G             -             U,A            C,C            U,A          A,G            G-bulge_Das
 IL   -             -             G,A            A,G            -            -              Tandem-GA
 IL   YCC           AAC           -              -              -            -              Twisted-up
 IL   YAA           RAN           -              -              -            -              UAA_GAN     # 23S rRNA, RNase P
 IL   RRRA          YYNR          -              -              -            -              J4a/4b      # riboswitch
 IL   AA            AA            -              -              -            -              J4/5-IL     # ribozyme
 IL   GUA           GG            -              -              -            -              GUA^GG_RRE
 IL   CAGG          AGCA          -              -              -            -              S_domain
 IL   GAAC          CUAG          -              -              -            -              Hook-turn
 IL   UGRAA         -             -              -              -            -              Docking-elbow
 IL   UU            AUU           -              -              -            -              pK-turn

#J3   S1(5'-3')   S2(5'-3')  S3(5'-3')                name
#------------------------------------------------------------------------------------------------------------------
J3      N         CUGA       A                        J3_hammerhead
J3      U         YUCUAC     AC                       J3_purine
J3      NN        NNNNNN     NN                       J3_typeA
J3      NNNN      NNNN       NNNN                     J3_typeB
J3      NN        NNN        NNNNNN                   J3_typeC
J3      -         UGAGA      -                        J3_TPP       #Breaker

#J4   S1(5'-3')   S2(5'-3')  S3(5'-3')   S4(5'-3')     name
#------------------------------------------------------------------------------------------------------------------
J4      -          AA         -           U            J4_HCV_IRES
J4      N          -          NNN         -            J4_tRNA
J4      N          N          NN          -            J4_manA
J4      R          -          -           R            J4_U1RNA

#BS  Loop(5'-3')  name
#------------------------------------------------------------------------------------------------------------------
BS       UKNRW            T-loop
BS       RRGU             LoopE-a
BS       RARR             LoopE-b
BS       AAYAARAACAANARR  CRC_binding
```

### 6 Inputs

#### The Stockholm format to describe a consensus structure

The input is a multiple sequence alignment in Stockholm format (https://en.wikipedia.org/wiki/Stockholm_format). The Stockholm format allows to provide a consensus structure for the alignment using the tag **#=GC SS_cons**. R-scape then can analyze the given consensus structure using option **-s**.

The Stockholm format uses symbols (), <>, {}, and [] to describe a nested structure. It also allows non-nested structures such as pseudoknots using the symbols Aa or Bb, ..., Zz. See file **tutorial/RF00162_SAM.pk.sto**, where a pseudoknot with 4 basepairs is annotated for the SAM-I riboswitch.

#### A extended Stockholm format to describe consensus structures with overlapping base pairs

The **#=GC SS_cons** annotation does not allow to display other pairs that overlap with the rest of the structure. These overlapping pairs can be base triplets, other non Watson-Crick interactions or even alternative structures. CaCo-Fold identifies a number of these overlapping pairs with covariation support. In order to annotate these overlapping basepairs, R-scape uses additional annotations in the form **#=GC SS_cons_1**, **#=GC SS_cons_2**,... See for example file **tutorial/RF00162_SAM.CaCoFold.sto**, where in addition to the pseudoknot there are two other overlapping pairs annotated as part of the structure.

You should use this augmented Stockholm format to input a consensus structure with overlapping basepairs. If you run

```
bin/R-scape -s tutorial/RF00162_SAM.CaCoFold.sto
```

you will see that the whole structure is taken into account in the statistical test.

### 7 Outputs

A Stockholm alignment file can include several different multiple sequence alignments (MSAs). For each alignment file rnafile.sto, R-scape produces the following output files, one for each individual alignment in an input Stockholm file:

**rnafile_msaname.cov** Tabular output with the significant pairs, with their score and E-value, estimated number of substitutions and power.

**rnafile_msaname.sorted.cov** Tabular output sorted from highest to lowest E-value.

**rnafile_msaname.power** Tabular output with the list of basepairs in the proposed RNA structure annotated their power. The file also reports the alignment power, and the expected number of basepairs to covary.

#### Covariation tabular output

The distribution includes in the directory tutorials/ examples of output files. If you run R-scape, the outputs will go into your current working directory (not necessarily tutorials/).

The output file **tutorial/updated_Arisong_1.cov** looks like this:

**> more tutorial/updated_Arisong_1.cov**

```
# Method Target_E-val [cov_min,cov_max] [FP | TP True Found | Sen PPV F]
# GTp    0.05           [-9.78,121.66]    [0 | 11 20 11 | 55.00 100.00 70.97]
#
# in_given  left_pos       right_pos        score          E-value       substitutions      power
#-------------------------------------------------------------------------------------------------------
*              94             110      57.27959        0.00760538      37             0.40
*              96             108      88.43400        0.000466924     26             0.28
...
```

The output file is a tabular list of significant pairs sorted by sequence positions:

`First column` indicates whether the significant pair is part of the given structure (*), or not. If the pair is not in the structure, we distinguish whether the pair is compatible with the given structure ($\sim$) or not (blank).

In addition, if the structure is provided by a PDB file (using the option `--pdb`), a non Watson-Crick/Watson-Crick base pair is designated by "**". A contact that is not a basepair is designated by: "$c\sim$" if compatible with all the basepairs, or by "$c$" otherwise.

`Second and third columns` are the two positions of the pair, $i \leq j$ respectively. Positions are relative to the input alignment.

`Fourth column` is the covariation score.

`Fifth column` is the E-value. Significant positions have E-values $<< 1$.

`Sixth column` is the estimated number of total substitutions in the two columns.

`Seventh column` is the basepair power or probability that it should covary.

The output file also includes two comment lines per alignment in the file:

`First comment line` describes properties of the alignment: number of sequence (nseq), alignment length (alen), average percentage identity (avgid), and number of base pairs (nbpairs). Values in parentheses correspond to the alignment as given. Values not in parentheses correspond to the analyzed alignment after the filters (for redundant sequences and gapped columns) have been applied.

`Second comment line` describes properties of the R-scape search: the covariation method (GTp), the E-value threshold (0.05), the range of scores for all pairs in the alignments (from -9.7 to 89.1), the number of covarying non base pairs (0), the number of covarying base pairs (11), the number of base pairs (20), and the total number of covarying pairs (11). Lastly we provide the sensitivity (SEN=55.00=11/20), positive predictive value (PPV=100.00=11/11), and F-measure (F=70.97 = 2 * SEN * PPV / (SEN+PPV)).

#### Power tabular output

The output file **tutorial/updated_Arisong_1.power** looks like this:

**> more tutorial/updated_Arisong_1.power**

```
# Power analysis of given structure
#
# covary  left_pos      right_pos    substitutions      power
#-------------------------------------------------------------
     *    94            110             37              0.40
          95            109             28              0.31
     *    96            108             26              0.28
     *    97            107             58              0.59
     *    98            106             45              0.48
     *    99            105             15              0.14
          100           104             20              0.21
.
.
.
     *    122           137             57              0.58
     *    123           135             72              0.68
     *    124           134             20              0.21
          125           133             31              0.34
#
# BPAIRS 20
# avg substitutions per BP  43.7
# BPAIRS expected to covary 8.3
# BPAIRS observed to covary 11
```

This file includes the list of all basepairs in the proposed structure given with the input alignment. Each basepair is annotated with the estimated number of substitutions and power.

#### Default graphical outputs

By default, the following files are also produced

**rnafile_msaname.R2R.sto** Stockholm file annotated by a modified version of the R2R program. This file includes the information necessary to draw the consensus structure, and to annotate the significantly covarying base pairs.

**rnafile_msaname.R2R.sto.{pdf,svg}** Drawing of the R-scape-annotated consensus secondary structure.

**rnafile_msaname.surv** A two column file with the survival functions (surv) for the covariation scores.

**rnafile_msaname.surv.ps** Plot of the score's survival function $P(X > \text{score})$. Drawing this file requires that program **gnuplot** is installed somewhere in the `${PATH}`, or that the environmental variable GNUPLOT pointing to a gnuplot executable is defined.

**rnafile_msaname.dplot.{ps,svg}** Dot plot of the consensus secondary structure annotated according to covariation. Drawing of this file requires that program **gnuplot** is installed somewhere in the `${PATH}`, or that the environmental variable GNUPLOT pointing to a gnuplot executable is defined.

For each alignment, **msaname** is given by <ACC>_<ID>, the combination of the accession #=GF AC <ACC> and name #=GF ID <ID> in the Stockholm-format markups (or one of two if the other in not defined). If none of those fields are defined, **msaname** is a number describing the order in the file of the given alignment.

#### Details about graphical outputs

Two files are produced per alignment in the input file:

File **tutorial/updated_Arisong_1.R2R.sto** is a Stockholm formatted alignment that includes the input alignment annotated with the consensus structure. This Stockholm file also includes the additional annotation required to use the drawing program R2R.

It is possible that the resulting drawing will show parts of the secondary structure occluded from each other (especially for long RNAs). Using this file, one can customize a different drawing of the structure using the R2R documentation, provided in lib/R2R/R2R-manual.pdf.

File **tutorial/updated_Arisong_1.R2R.sto.pdf** depicts the consensus structure with the R-scape covariation annotation using the R2R software to depict the alignment. The defaults used by R-scape to depict the alignment positions are given in this legend,

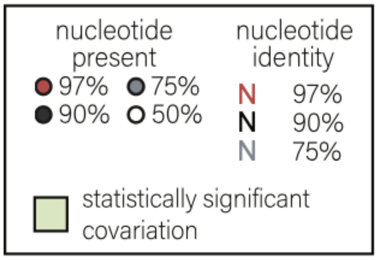

File **tutorial/updated_Arisong_1.surv** looks like this:

**> more tutorial/updated_Arisong.surv**

```
121.795428      0.05
95.862635       0.1
89.113004       0.15
...
 &
63.890698       0.000485437
58.917286       0.000970874
47.904730       0.00145631
...
 &
81.652885       2.40385e-06
77.745204       4.80769e-06
77.034717       7.21154e-06
...
 &
256.788050      2.64342e-17
256.432807      2.7899e-17
256.077563      2.94449e-17
...
 &
```

The first column is a covariation score (x). The second column is the survival function $P(X > x)$, that is the frequency of pairs having score larger than x. The file includes four survival functions separated by a "&" line. The three survival functions correspond to:

First functions: the given alignment, proposed base pairs. (This section is empty if no secondary structure is proposed.)

`Second functions:` the given alignment, not proposed pairs.

`Third function:` the aggregation of all null alignments, all possible pairs.

`Fourth function:` the expected null survival function according to the tail Gamma fit.

#### Using option –cacofold

If the option `--cacofold` is used, R-scape produces the following additional files describing the maximal-covariation optimal secondary structure:

**rnafile_msaname.fold.sto** The original alignment with the R-scape structural annotation

**rnafile_msaname.fold.R2R.sto** File used by R2R to display the R-scape structure

**rnafile_msaname.fold.R2R.sto.{pdf,svg}**

**rnafile_msaname.fold.surv**

**rnafile_msaname.fold.surv.{ps.svg}**

**rnafile_msaname.fold.dplot.{ps,svg}**

These files are formatted identically to those describing the given consensus structure.

#### Graphical outputs per alignment

Three plots are produced per alignment in the input file:

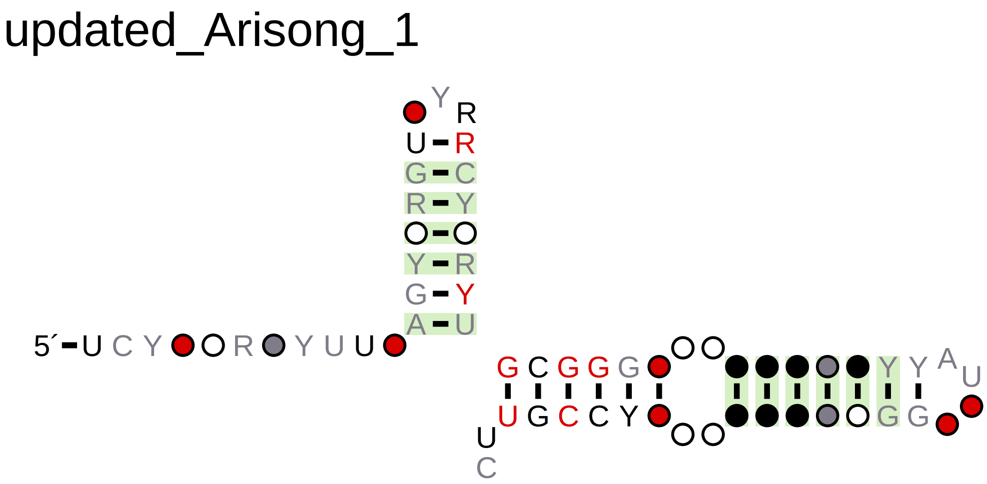

Figure 2: **`tutorial/updated_Arisong_1.R2R.sto.`{`pdf,svg`}`:` annotated consensus secondary structure.** Base pairs with covariation scores equal or below the target E-value (0.05 as default) are depicted in green. By default only positions in the alignment with more than 50% occupancy are depicted (unless they form a base pair). Option `--r2rall` forces the depiction of all positions in the alignment.

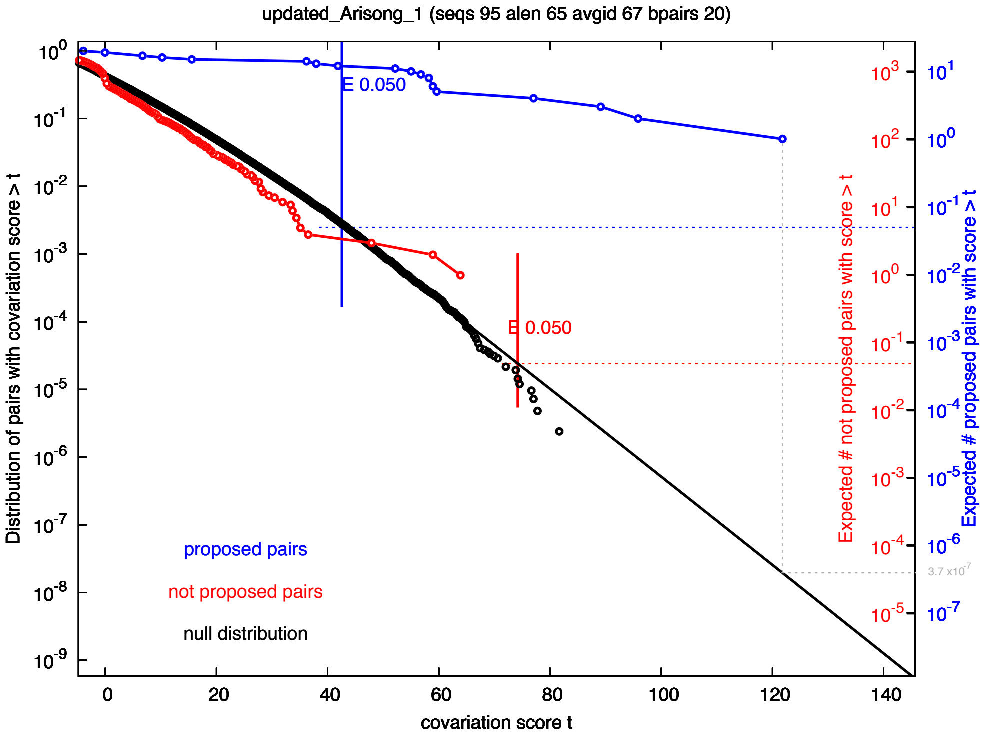

Figure 3: **tutorial/updated_Arisong_1.surv.{ps,svg}: covariation scores survival function** $P(X > x)$**.** The survival function of scores for all pairs in the given alignment is depicted in blue. The survival function for the null alignments is depicted in black. A black line indicates to fit to a truncated Gamma distribution of the tail of the null distribution. In red, we plot the survival function of scores for the pairs in the given alignment excluding those proposed as base pairs. For a particular pair, as an example the highest scoring one from the distribution of proposed pairs (blue), we obtain its E-value by drawing a vertical (gray) line from the point to the null distribution (black). The corresponding value in the blue scale gives us the E-value for that pair (in this example, $3.7 \cdot 10^{-7}$).

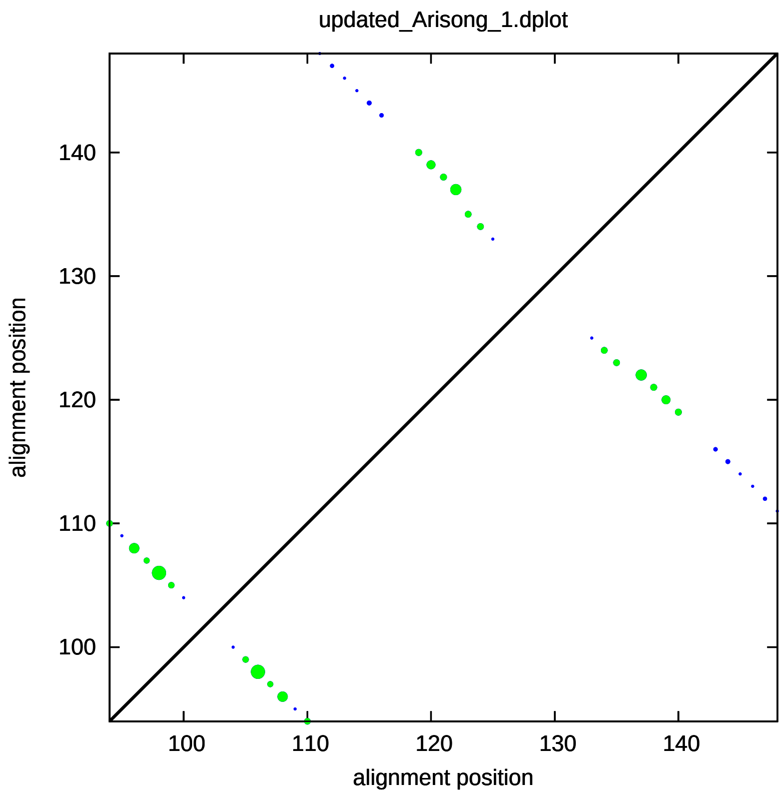

Figure 4: **`tutorial/updated_Arisong_1.dplot.`{`ps,svg`}`:` dotplot.** Dot size is proportional to the covariation score. In blue we depict the consensus base pairs; in green, the consensus base pairs that show significant covariation; in orange (none shown in this plot), we depict other pairs that have significant covariation, are not part of the consensus secondary structure but are compatible with it; in black we depict other significant pairs. Position are relative to the original input alignment (before any gapped column is removed).

### 8 Options

The whole list of options can be found using

**> R-scape -h**

Some important options are:

#### Covariation statistic options

**-E <x>**

Target E-value is $x \geq 0$.

**--GT, --MI, --MIr, --MIg, --CHI, --OMES, --RAF, --RAFS,**

We favor the G-test covariation statistic, but a total of eight covariation statistics are currently implemented in R-scape. For each covariation statistic (GT, for instance), R-scape can also calculate its average product correction (GTp) and its average sum corrections (GTa). For each option above, appending "p" or "a" chooses one of the corrections. For example, `--GT` does the G-test statistic, `--GTp` does the APC-corrected G-test statistic, `--GTa` does the ASC-corrected G-test statistic.

The R-scape default is `--GTp`.

Details of the definition and provenance of the different covariation statistics can be found in the R-scape manuscript: Rivas, E. & Eddy S. E., *"A statistical test for conserved RNA structure shows lack of evidence for structure in lncRNAs"*.
In a nutshell, given two alignment columns $i, j$,

| | | | |
|---|---|---|---|
| G-test:(Woolf, 1957) | $\mathrm{GT}(i,j)$ | $=$ | $2\sum_{a,b} \mathrm{Obs}_{ij}^{ab} \log \frac{\mathrm{Obs}_{ij}^{ab}}{\mathrm{Exp}_{ij}^{ab}}$, |
| Pearson's chi-square: | $\mathrm{CHI}(i,j)$ | $=$ | $\sum_{a,b} \frac{\left(\mathrm{Obs}_{ij}^{ab}-\mathrm{Exp}_{ij}^{ab}\right)^2}{\mathrm{Exp}_{ij}^{ab}}$, |
| Mutual information:(Shannon, 1948; Gutell et al., 1994) | $\mathrm{MI}(i,j)$ | $=$ | $\sum_{a,b} P_{ij}^{ab} \log \frac{P_{ij}^{ab}}{p_i^a\, p_j^b}$, |
| MI normalized:(Martin et al., 2005) | $\mathrm{MIr}(i,j)$ | $=$ | $\frac{\mathrm{MI}(i,j)}{H(i,j)} = \frac{\mathrm{MI}(i,j)}{-\sum_{a,b} P_{ij}^{ab} \log P_{ij}^{ab}}$, |
| MI with gap penalty:(Lindgreen et al., 2006) | $\mathrm{MIg}(i,j)$ | $=$ | $\mathrm{MI}(i,j) - \frac{N_{ij}^G}{N}$, |
| Obs-Minus-Exp-Squared:(Fodor and Aldrich, 2004) | $\mathrm{OMES}(i,j)$ | $=$ | $\sum_{a,b} \frac{\left(\mathrm{Obs}_{ij}^{ab}-\mathrm{Exp}_{ij}^{ab}\right)^2}{N_{ij}}$, |
| RNAalifold (RAF):(Hofacker et al., 2002) | $\mathrm{RAF}(i,j)$ | $=$ | $\mathrm{B}_{i,j}$, |
| RNAalifold Stacking (RAFS):(Lindgreen et al., 2006) | $\mathrm{RAFS}(i,j)$ | $=$ | $\frac{1}{4}\left(\mathrm{B}_{i-1,j+1} + 2\,\mathrm{B}_{i,j} + \mathrm{B}_{i+1,j-1}\right)$. |

where $a, b$ are (non-gap) residues; $N$ is the total number of aligned sequences; $\mathrm{Obs}_{ij}^{ab}$ is the observed count of $a : b$ pairs in columns $i, j$ (only counting when both a,b are residues); $N_{ij}$ is the total number of residue pairs in columns $i, j$ (only counting when both a,b are residues); $P_{ij}^{ab}$ is the observed frequency of pair $a : b$ in columns $i, j$ ($P_{ij}^{ab} = \frac{Obs_{ij}^{ab}}{N_{ij}}$); $\mathrm{Exp}_{ij}^{ab} = N_{ij} p_i^a p_j^b$ is the expected frequency of pair $a : b$ assuming $i, j$ are independent, where $p_i^a$ are the marginal frequencies of $a$ residues in column $i$ (averaged to all other positions) ($p_i^a = \frac{1}{L-1}\sum_{j \neq i}\sum_b P_{ij}^{ab}$); $N_{ij}^G = N - N_{ij}$ is the number of pairs involving at least one gap symbol; the definition of $\mathrm{B}_{i,j}$ used in the RAF and RAFS statistics is involved, a concise definition can be found elsewhere (Lindgreen et al., 2006).

The background corrections (Dunn et al., 2007) for a given covariation statistic above $\mathrm{COV}(i,j)$ are,

| | | | |
|---|---|---|---|
| Average product correction | $\mathrm{COVp}(i,j)$ | $=$ | $\mathrm{COV}(i,j) - \frac{\mathrm{COV}(i)\mathrm{COV}(j)}{\mathrm{COV}}$, |
| Average sum correction | $\mathrm{COVa}(i,j)$ | $=$ | $\mathrm{COV}(i,j) - \left(\mathrm{COV}(i) + \mathrm{COV}(j) - \mathrm{COV}\right)$. |

**`--C2, --C16, --CWC`**

For all the covariation statistics (except RAF and RAFS), one can do a 16-component (C16) or a two-component (C2) calculation, depending on whether it uses the 16 possible pair combinations, or those are group in two classes depending on whether they form a Watson-Crick pair (6 cases, including U:G and G:U), or whether they do not (10 cases).

R-scape's default is the 16 component covariation statistic, unless the number of sequences in the alignment is $\leq$ 8 or the length of the alignment is $\leq$ 50, in which case it uses the two-class covariation statistic.

Option CWC (implemented for GT, GTp and GTa) uses a 6 component covariation statistic which only considers Watson-Crick-like pairs: A-U, U-A, C-G, G-C, G-U, U-G.

#### Options to calculate power

**`--singlesubs`**

Default option. The base pair substitutions are calculated as the sum of the individual substitutions observed for each of the positions in the base pair.

**`--joinsubs`**

The base pair substitutions are calculated as the sum of the individual substitution observed for each positions in the base pair, but using only sequences in which both positions are occupied, that is, cases where one of the two positions is a gap are ignored.

**`--doublesubs`**

The base pair substitutions are calculated a the subset of substations in which both residues have changed.

#### Covariation aggregation options

**`--fisher,--sidak`**

Two different options to produce aggregated E-values calculated for each helix in the proposed or CaCoFold structure.

**`--lancaster,--wfisther`**

Two different options to produce aggregated E-values calculated for each helix in the proposed or CaCoFold structure.

The Lancaster aggregation uses the number of substitutions per base pair. The weighted fisher (wfisher) aggregation uses the power per base pair. These two options require the use of the default option **`--singlesubs`** in order to calculate the number of substitutions and power for each base pair respectively.

**`--lancaster_join,--wfisther_join`**

These two options require the use of the option **`--joinsubs`** in order to calculate the number of substitutions and power for each base pair respectively.

**`--lancaster_double,--wfisther_double`**

These two options require the use of the option **`--doublesubs`** in order to calculate the number of substitutions and power for each base pair respectively.

#### Search options

**-s**

The “two-set test” option. This option requires that a structure is provided with the alignment. If option `-s` is used, R-scape performs two independent test, one for the given structure, another for all other possible pairs. The default is a “one-set test” in which all possible pairs in the alignment are tested equivalently.

**--cacofold**

A CaCoFold structure is computed that includes all significant base pairs. All files related to this CaCoFold structure include the suffix `.cacofold.`

When option `--cacofold` is used, a file with the original alignment annotated with the R-scape structure in Stockholm format is produced. This alignment has the suffix `.cacofold.sto`.

**--naive**

Reports the laundry list of all covariation scores, without any statistical significance (E-value) associated to them. No null alignments are created.

**--tstart <n>**

Analyze starting from position $n >= 1$ in the alignment.

**--tend <n>**

Analyze ending at position $n <= L$ in the alignment.

**--window <n>**

R-scape can be run in a window scanning version for long alignments. The window size is $n > 0$.

**--slide <n>**

In scanning mode, this options sets the number of positions to move from window to window, $n > 0$.

**--vshuffle**

Vertical shuffle, a developers tool. Before performing any analysis, it shuffles all residues in each alignment column independently.

**--cshuffle**

Column shuffle, a developers tool. Before performing any analysis, it shuffles all columns in the alignment.

**--givennull <f>**

Use histogram provided in file $< f >$ as null.

#### Input alignment options

**-I <x>**

Only sequences with less than $0 < x \leq 1$ pairwise similarity are considered in the analysis. Pairwise % identity is defined as the ratio of identical positions divided by the minimum length of the two sequences. If this option is not used all (weighted) sequences are used in the analysis.

**--gapthresh <x>**

Only columns with less than $0 < x \leq 1$ fraction of gaps are considered in the analysis.

**--consensus**

If the alignment has a GC "seq_cons" field, only consensus positions will be analyzed.

**--submsa <n>**

Analyzes a random subset of the input alignment.

**--treefile <f>**

A phylogenetic tree in Newick format can be given (by default a tree is created from the alignment using the program FastTree (v2.1.11) (Price et al., 2010)). R-scape checks that the number of taxa and the names of the taxa matches for all alignments analyzed.

**--ntree <n>**

Number of different FastTree trees to use when generating the null alignments. Default is one tree, the one resulting from feeding the input alignment to FastTree. if `--ntree` > 1, the rest of the trees are generated after randomly rearranging the sequences in the alignment.
Because FastTree is not deterministic, altering the order of the sequences in the alignment can result in slightly different trees, and in some rare occasions that can results in different distribution of null covariation scores. In those cases, it is recommended to generate null alignments from different trees obtained from randomly rearrange the sequences in the alignment.
Option `--ntree <n>` is incompatible with option `--treefile <f>` which inputs a particular tree.

#### Options for producing a CaCoFold structure

**--cacofold**

When using the option `--cacofold`, R-scape engages the CaCoFold algorithm to produce a predicted structure. The CaCoFold algorithm incorporates all positive (significantly covarying) base pairs, and prevents any negative pair (pairs that have power of covariation but not covariation) from happening. The CaCoFold algorithm uses a recursive cascade of constrained foldings. The first fold uses the RBG probabilistic grammar, the rest use the G6X probabilistic grammar. To find secondary structure and RNA 3D motifs:

**--cacofold --r3d**

to find the secondary structure and RNA 3D motifs.

**--r3dfile <f>**

provide a customized r3d file. The defaults file is at `data/r3d/R3D.grm`
Regarding the predicted structure, CaCoFold can use one two algorithms:

**--cyk**

Default option. Each folding reports the structure with the best probability using the CYK algorithm.

**--decoding**

This options returns the structure obtained by posterior decoding.
Posterior decoding usually performs better than CYK. Both algorithm has the same algorithmic complexity. CYK is faster.
Several additional options can be used in combination with `--cacofold`,

**--refseq**

By default the CaCoFold algorithm folds a profile sequence built from the alignment. Using this option, the sequence to fold is a consensus reference sequence.

**--E_neg <x>**

Pairs with E-value larger than the E-value cutoff but smaller than ¡x¿ will not be called negatives regardless of their covariation power. Default for E_neg is 1.0.

**--lastfold**

This option forces one last alternative fold (using grammar G6X) after all covarying base pairs have already been integrated into the structure. By default this last fold is not performed. In the absence of any covarying base pair, one fold is performed using grammar RBG.

**--show_hoverlap**

This option leaves the alternative helices unmodified. By default, alternative structures are trimmed down to show no overlap with helices from the previous layers.

**--covmin <n>**

Minimum distance between position to report significant covariation. Default is 1, which means that significant covariation between contiguous positions are reported.

**--allow_negatives**

This option (just for developers) allows all base pairs to form regardless of their power.

**--Rfam**

This option is meant to be used by the Rfam curators when using CaCoFold to propose improved consensus structures for an Rfam family. It removes covariation that cannot be taking into account by the Rfam models. Thus, maybe missing important covariation information that is not compatible with RNA secondary structure.

**--rmcoding**

This option removes from the R2R display the covariation that appears to be due to protein-coding codon signals instead of RNA structure. This affect only to the display figures not to anything in the analysis of covariation.
Using options `--cacofold --Rfam`, the CaCoFold structure is trimmed such that:

- Base pairs have to have at least 3 nucleotides of separation (covarying pairs removed if they don't).

- Overlaps between helices are trimmed down if possible without removing any covarying pair.
- Pseudoknots (pk) are kept, but alternative motifs identified as: triplets (tr), cross (xcov), or side (scov) covariations are removed.
- Base pairs which appear to be non WC (defined by the observed frequency in the alignment of the pair being A:U, U:A, C:G, G:C, G:U or U:G being less than 0.3) are removed, even if they covary.

#### Options for importing a structure

R-scape does not require to input a structure (either a RNA structure or a protein contact map). By default R-scape analyzes all possible pairs in the alignment.

There are two ways to provide a contact map (or structure):

- By providing the alignment in Stockholm format with a "ss_cons" field including the consensus structure for the alignment. (For RNA alignments only.)
- By analyzing a 3D structure provided in a PDB file. (For either RNA or peptide alignments.)

These two methods can be combined together. For a nucleotide alignment, if both a consensus structure is present in the alignment, and a PDB file is provided (using option `--pdb`), the consensus structure will be extended by the information provided by the pdbfile. To ignore the consensus structure use option `--onlypdb`.

From the PDB file we obtain three types of structural pairs:

- **Contacts:** defined as those two residues at a close spatial distance (specified by the user with option `--cntmaxD`).
- **Basepair:** RNA base pairs.
  RNA Basepair are calculated using the program `rnaview` (Yang et al., 2003).

  These RNA base pairs can be further classified in two types:

  - **Watson-Crick base pairs:** the canonical RNA base pairs. mostly A:U, G:C, or G:U pairs. (H-bond interactions between two W-C faces in cis).
  - **Other base pairs:** the non-canonical RNA base pairs (all other types of H-bond interactions, 12 different types).

  Contacts and RNA base pairs are extracted as follows:

- The spatial distance between any two residues is calculated as the minimal Euclidean distance between any two atoms (excluding H atoms). Any two pairs at a distance not larger than a maximum value (`contmaxD`) are called a "contact".
- RNA base pairs are obtained using the program `rnaview` (Yang et al., 2003)
  (http://ndbserver.rutgers.edu/ndbmodule/services/download/rnaview.html).
  The RNA base pair annotation takes precedent over the annotation as "contact".

The options that control the input of a structure or contact map are:

**`--pdb <s>`**

Reads a pdbfile associated to the alignment, and extracts the contacts from it.
A ".cmap" file is produced reporting the structure obtained from the PDB file.
Option `--pdb` is incompatible with `--cacofold`.

**--cntmaxD <x>**

Maximum distance (in Angstroms) allowed between two residues to define a "contact" is $\langle x \rangle$.

**--cntmind <n>**

Minimum distance (in residue positions) in the backbone between two residues required to define a "contact" is $\langle n \rangle$.

**--onlypdb**

Reads the structure from the pdbfile and ignores the alignment consensus structure (if provided).

**--draw_nonWC**

Adds the non-canonical base pairs into the structure graphical output. For clarity, the default is to draw only the Watson-Crick base pairs. This option affects only the drawing of the structure. All base pairs (canonical or not) are used as part of the structure to perform the two-set statistical test.

Example of reading a structure from a PDB file for the FMN riboswitch:

**> bin/R-scape --cntmaxD 4 --cntmind 3 --pdb tutorial/3f2q.pdb -s --onlypdb tutorial/RF00050.sto**

This command line extracts contacts from the pdb file that are at a Euclidean distance $\leq 4\AA$ in the PDB structure, and such that they are at least 3 residues apart in the backbone.
The output is

```
# R-scape :: RNA Structural Covariation Above Phylogenetic Expectation
# R-scape 0.8.1 (Jul 2018)
# Copyright (C) 2016 Howard Hughes Medical Institute.
# Freely distributed under the GNU General Public License (GPLv3).
# - - - - - - - - - - - - - - - - - - - - - - - - - - - - - - - - - - - - -
# Two-set statistical test (one test for annotated base pairs, another for all other pairs)
#
# Structure obtained from the pdbfile
# ij in alignment | ij in pdbsequence | basepair type
# 3 218 | 1 112 | WWc
# 4 216 | 2 110 | CONTACT
# 4 217 | 2 111 | WWc
# 4 218 | 2 112 | CONTACT
# 5 216 | 3 110 | WWc
# 5 217 | 3 111 | CONTACT
# 6 215 | 4 109 | WWc
# 6 216 | 4 110 | CONTACT
# .
# .
# .
# 192 202 | 87 96 | WWc
# 192 203 | 87 97 | CONTACT
# 193 198 | 88 92 | CONTACT
# 193 201 | 88 95 | WWc
# 193 202 | 88 96 | CONTACT
# 195 197 | 89 91 | CONTACT
# 195 198 | 89 92 | WHt
# 198 200 | 92 94 | CONTACT
# 198 201 | 92 95 | CONTACT
# 205 207 | 99 101 | CONTACT
# PDB:      versions/rscape/rscape_v0.8/tutorial/3f2q.pdb
# contacts  169 (49 bpairs 35 wc bpairs)
# maxD      4.00
# mind      3
# distance  MIN
# L         139
# alen      221
# pdblen    112
# ::[[[[[[[[,,,,,,<<<_______>>>,((((<<<<_________________AA>>>>,,<<<----------<____>>>>,,,<<<<<_______>>>>>aa))AAAA----))a
```

```
# MSA RF00050_FMN.3f2q nseq 144 (144) alen 139 (221) avgid 69.18 (68.15) nbpairs 49 (0)
#
# Method Target_E-val [cov_min,conv_max] [FP | TP True Found | Sen PPV F]
# GTp    0.05         [-9.78,216.11]     [1 | 14 49 15 | 28.57 93.33 43.75]
#
#       left_pos       right_pos        score           E-value
#------------------------------------------------------------------
*            171            183       216.11095        1.6421e-10
*            170            184       211.69081        2.76699e-10
*            192            202       168.72417        4.95548e-08
*              8            213       149.71776        4.89982e-07
*            172            182       138.66664        1.84675e-06
*            169            185       137.23189        2.21548e-06
**            16             30        133.44999        3.53772e-06
*              5            216       131.02575        4.70876e-06
*             84            186       125.60806        9.0169e-06
*             17             29       112.04610        4.62895e-05
*              7            214       111.12654        5.13519e-05
*              6            215       96.43781        0.00029929
*             36             87       96.32752        0.00029929
*             94            163       78.81578        0.0024303
               7            213       107.68588        0.0147937
```

All coordinates are relative to the input alignment. The annotation of all types of RNA base pairs (WWc, WWt, WHc,...) is produced by the program `rnaview` (Yang et al., 2003).

#### Options for type of pairs tested

When performing the two-class statistical test (option `-s`) using a pdbfile to read the structure, there are different options as to which types of base pairs are used to define the sample size for the base pairs test.

The options are:

**--samplecontacts**

The basepair statistical test includes all the contacts identified in a PDB or/and as a RNA secondary structure included with a input alignment in Stockholm format. This is the default option for amino acid alignments if a PDB file is provided.

**--samplebp**

For RNA alignments with only. The basepair statistical test includes base pairs of all 12 possible types. This is the default option for RNA/DNA alignments if a PDB file is provided.

**--samplewc**

For RNA alignments only. The basepair statistical test includes only the canonical (Watson-Crick/Watson-Crick type) base pairs (A:U, G:C, G:U). This is the default option for RNA/DNA alignments if a consensus secondary structure is provided.

#### Output options

**--roc**

Produces a tabular output that provides statistics for each score value.

File **tutorial/updated_Arisong.roc** looks like:

**> more tutorial/updated_Arisong.roc**

```
# MSA nseq 95 alen 65 avgid 66.352419 nbpairs 20 (20)
# Method: GTp
#cov_score  FP  TP Found  True  Negatives  Sen   PPV     F       E-value
121.79543   0   2  2      20    2060       10.00 100.00  18.18   4.07104e-05
121.44018   0   2  2      20    2060       10.00 100.00  18.18   4.29443e-05
```

```
121.08494   0   2  2      20    2060       10.00 100.00  18.18   4.53006e-05
120.72970   0   2  2      20    2060       10.00 100.00  18.18   4.53006e-05
...
```

This file produces a tabular output for each alignment as a function of the covariation score, for plotting ROC curves. The values in the file are described by the comment line. Notice that the number of Trues (column 5) and Negatives (column 6) are fixed for a given secondary structure and do not change.

**--outmsa <f>**

The actual alignment analyzed can be saved in Stockholm format to file <f>.

**--outtree <f>**

The phylogenetic tree (created using the program FastTree) can be saved in Newick format to file <f>.

**--savenull**

Saves a histogram with the null distribution to file rnafile msaname.null.

#### Plotting options

**--nofigures**

None of the graphical outputs are produced using this option.

**--r2rall**

Forces R2R to draw all positions in the alignment. By default only those that are more than 50% occupied or are base paired are depicted.

#### Other options

**--seed <n>**

Sets the seed of the random number generator to <n>. Use n = 0 for a random seed.

### 9 Some other topics

#### How do I cite R-scape?

Rivas, E. *et al.*, *"A statistical test for conserved RNA structure shows lack of evidence for structure in lncRNAs"*, Nature Methods 14, 4548 (2017).
https://www.nature.com/articles/nmeth.4066

Rivas, E. and Eddy, S. E., *"Response to Tavares et al., Covariation analysis with improved parameters reveals conservation in lncRNA structures"*, (2018).
https://doi.org/10.1101/2020.02.18.955047.

Rivas, E. *et al.*, *"Estimating the power of sequence covariation analysis for detecting conserved RNA structure"*, *Bionformatics*, 36, 30723076, (2020).
https://doi.org/10.1093/bioinformatics/btaa080

Rivas, E., *"RNA covariation at helix-level resolution for the identification of evolutionarily conserved RNA structure"*, *PLOS Comput Biol, 19(7): e1011262, (2023).*
https://doi.org/10.1371/journal.pcbi.1011262.

#### How do I cite CaCoFold?

Rivas, E., *"RNA structure prediction using positive and negative evolutionary information"*, *PLOS Comput Biol*, 16(10), e1008387, (2020).
https://doi.org/10.1371/journal.pcbi.1008387.

#### How do I cite CaCoFold-R3D?

Karan, A and Rivas, E., *"All-at-once RNA folding with 3D motif prediction framed by evolutionary information"*, *bioRxiv*, Dec (2024).
https://doi.org/10.1101/2024.12.17.628809, 20 Dec 2024.

You should also cite what version of the software you used. We archive all old versions, so anyone should be able to obtain the version you used, when exact reproducibility of an analysis is an issue.

The version number is in the header of most output files. To see it quickly, do something like `R-scape -h` to get a help page, and the header will say:

```
# R-scape :: RNA Structural Covariation Above Phylogenetic Expectation
# R-scape 2.5.7 (March 2025)
# Copyright (C) 2017-2025 Elena Rivas, Harvard University.
# Freely distributed under the GNU General Public License (GPLv3).
# - - - - - - - - - - - - - - - - - - - - - - - - - - - - - - - - - - - - - - -
```

So (from the second line there) this is from R-scape v2.5.7.

#### How do I report a bug?

Before we can see what needs fixing, we almost always need to reproduce a bug on one of our machines. This means we want to have a small, reproducible test case that shows us the failure you're seeing. So if you're reporting a bug, please send us:

- A brief description of what went wrong.

- The command line(s) that reproduce the problem.
- Copies of any files we need to run those command lines.
- Information about what kind of hardware you're on, what operating system, and what compiler and version you used, with what configuration arguments.

### 10 Acknowledgments

We thank S.E. Roian Egnor for suggesting the name R-scape, and the Centro de Ciencias de Benasque Pedro Pascual in Spain, for their hospitality, over numerous and wonderful summers.
